# Activity-Dependent Localization and Heterogeneous Dynamics of STIM1 and STIM2 at ER-PM contacts in Hippocampal Neurons

**DOI:** 10.1101/2024.12.23.630200

**Authors:** Arun Chhikara, Filip Maciąg, Nasrin Sorush, Martin Heine

## Abstract

Stromal interaction molecules (STIMs) are calcium sensors integral to store-operated calcium entry (SOCE), a process critical for non-excitable cells and contributing to homeostatic functions in neurons. Upon depletion of Ca^2+^ from the endoplasmic reticulum (ER), STIMs translocate to ER-plasma membrane (PM) junctions to contact the inner leaflet of the plasma membrane. Using single-particle tracking (SPT), we characterized the dynamic properties of neuronal STIM1 and STIM2 in hippocampal neurons.Our data reveal that STIMs exhibit heterogenous dynamics in dendrites and axons, while only transiently visiting synaptic compartments. A substantial fraction of STIM2 proteins define ER-PM contacts under resting conditions, whereas STIM1 proteins are recruited to ER-PM junctions during strong activation of glutamatergic synapses. Junctions organized by K_V_2.1 channels are not particularly enriched with STIM proteins. Activity-dependent confinement of STIM proteins is not influenced by L-type calcium channel (Ca_V_1.2) activity. We propose that STIM proteins predominantly regulate the contact area and frequency of contacts between ER and PM.

## 1. Introduction

ER-PM contacts are specialized regions where the endoplasmic reticulum (ER) and plasma membrane (PM) come into close proximity (10 nm in neurons) facilitating interactions between proteins in both membranes (Orci et al., 2009). STIM proteins (STIM1 and STIM2 isoforms in mammals) are calcium sensing proteins in the ER that can directly interact with PM proteins at the ER-PM junction (Ambudkar et al., 2017; Wang et al., 2010; Yuan et al., 2009). In non-neuronal cells, the interaction between STIM and Orai proteins opens calcium-release activated channels (CRACs), which are the major component of the store-operated calcium entry (SOCE) (Putney et al., 2001; Voelkers et al., 2010). This interaction is triggered by depletion of calcium stores in the ER, which is sensed by STIM proteins, and is an essential mechanism to control intracellular calcium signaling in non-excitable cells (Gross et al., 2007; Liou et al., 2005; Roos et al., 2005). In polarized cells like neurons, excitability and local communication are highly dependent on the distribution of voltage-gated channels (VGCCs) and local calcium signaling. Despite the abundance of STIM proteins in the nervous system (Dziadek & Johnstone, 2007; Gross et al., 2007; Klejman et al., 2009), the diversity of voltage and ligand-gated calcium channels in neurons raises questions about the relative relevance of SOCE in neuronal function (Lu & Fivaz, 2016). Neuronal ER-PM junctions cover approximately 10-20% of the total PM area, which is in stark contrast to the mere 1-2% in non-excitable cells (Giordano et al., 2013; Orci et al., 2009). Neuronal ER-PM junctions are especially prominent in the soma, but the high abundance of ER-membrane structures in postsynaptic spines suggests that ER-PM junctions may also be present within the synaptic compartment. About 40% of dendritic spines are visited by ER-membrane structures, whereas contacts within the presynapse are not supported by electron microscopy-based reconstructions (Perez-Alvarez et al., 2020; Y. Wu et al., 2017).

Despite the paucity of structural studies which suggest that the STIM-Orai pair functions as an additional calcium conduction unit, genetic manipulations of STIMs in neurons suggest that STIMs act as regulators of synaptic function and connectivity, potentially through transient ER-PM junctions along neurites. (Lalonde et al., 2014; Serwach & Gruszczynska-Biegala, 2020). Intracellular Ca^2+^ stores are known to impact neurotransmitter release through calcium-induced calcium release (CICR) (Emptage et al., 2001), postsynaptic calcium signaling (Dittmer et al., 2017; Dittmer & Dell’Acqua, 2024) and to influence the stabilization and formation of spines (Basnayake et al., 2021; Perez-Alvarez et al., 2020). Spatially restricted synaptic stimulation along dendrites supports the idea of local recruitment of STIM proteins (Dittmer et al., 2017). In addition to ORAI channels, other interaction partners, particularly in somatic and dendritic membranes such as AMPA and NMDA receptors, are proposed to be modulated by the activity-dependent recruitment of STIM proteins (Garcia-Alvarez, Shetty, et al., 2015; Gruszczynska-Biegala et al., 2016, 2020; Maneshi et al., 2020; Serwach et al., 2023; Yap et al., 2017; reviewed in Maciąg et al., 2024). STIM proteins are also suggested to directly interact and modulate VGCCs at ER-PM contacts, mediated by NMDA receptor-induced recruitment of STIM proteins near activated synapses on the dendrite (Dittmer et al., 2017; Park et al., 2010; Wang et al., 2010). The conductivity of STIM-ORAI dependent CRAC channels is much lower than known for VGCCs (Rothberg et al., 2013), suggesting a local rather than a global impact of such contacts.

As extensively investigated in lymphocytes and other non-excitable cells, the diffusive organization of STIM proteins is instructive to understand their function (Gil et al., 2021; Prakriya & Lewis, 2015). STIM proteins are dynamic; their activation and interactions with other proteins result in a diffusion trap at ER-PM junctions (M. M. Wu et al., 2014). Such dynamic STIM interactions, resulting in transient ER-PM contacts, position STIM-enriched ER-PM junctions as attractive modules for synaptic plasticity. These transient STIM-dependent junctions link synaptic calcium influx through VGCCs with ER-resident ryanodine receptors (Dittmer et al., 2017). However, structural evidence that STIM-ORAI contacts indeed predominate ER-PM junctions in synapses is very limited. In order to understand the dynamic nature of STIM mediated contacts in neurons from a molecular dynamics perspective, we employed super-resolution single-particle tracking (SPT).

We focused on the predominant splice variant of both STIM1 and STIM2, as these proteins exhibit widespread expression in the brain with regional and cellular heterogeneity (Ryazantseva et al., 2018; Skibinska-Kijek et al., 2009; Zhang et al., 2014). Despite sharing homology, STIM1 and STIM2 display distinct molecular mechanisms and functional kinetics. These subtle differences have been implicated in divergent phenotypic outcomes, with calcium binding affinity often cited as a key determinant of isoform-specific effects (Chin-Smith et al., 2014; Emrich et al., 2021).

We characterized the diffusion properties of STIM proteins in axons and dendrites. STIM proteins are mobile throughout the entire neuron; however, characteristic differences exist between STIM isoforms and neuronal compartments. At rest, both isoforms are highly diffusive in axons. In dendrites, STIM2 is pre-clustered in the basal state, while the more highly diffusive STIM1 forms clusters only after ER Ca^2+^ store depletion. Store depletion failed to induce cluster formation of either isoform in axons. In synaptic compartments, STIM proteins did not predominantly cluster in the presynaptic active zone or in the proximity of the postsynaptic density (PSD), indicating a segregation of calcium signaling within and around the synapse. NMDA receptor activation strongly stabilized STIM proteins, independent of CaV1.2 channel activity, and was much more potent than passive ER Ca^2+^ store depletion. We further observed that stabilized STIM proteins localize at ER-PM junctions, distinct from those populated by KV2.1.

## 2. Results

### 2.1. STIM protein dynamics differs between dendrites and axons

STIM1 and STIM2 (human isoforms denoted in uppercase, mouse isoforms as ‘Stim’) both undergo alternative splicing, resulting in variants with distinct functions (Miederer et al., 2015; Ramesh et al., 2021; Rosado et al., 2016). First, we identified the predominant splice variants of Stim proteins across brain regions (cortex, hippocampus, cerebellum) using the 2ΔCt qRT-PCR method (**Fig. Sup. 1**). Notably, Stim2 expression was higher than Stim1 in the hippocampus and cerebellum, and Stim1s (NM_009287.5) and Stim2.2 (NM_001081103.2) were the most abundant variants, convergent with other studies (Gruszczynska-Biegala et al., 2011). Based on sequence similarities and abundance in hippocampal neurons, human STIM1S (NM_003156.4) and STIM2.2 (NM_020860.4) were chosen as expression constructs. The subcellular distribution of STIM proteins in cultured hippocampal neurons was examined using three distinct experimental strategies to ensure accurate interpretation of their dynamic behavior: overexpression, re-expression, and CRISPR-Cas9-mediated endogenous tagging. All three approaches utilized N-terminal Halo-tagging driven by a human synapsin promoter and integrated into recombinant adeno associated vector (rAAV), allowing real-time tracking of individual STIM proteins in living neurons using membrane-permeable Halo-ligands conjugated to the fluorophore JF646. These constructs are referred to as STIM1 and STIM2 unless specified otherwise. The initial functionality of the constructs was tested in HEK293T cells (**Fig. Sup. 2**). On DIV 7 neurons were co-transduced with pre- or postsynaptic markers (Synaptophysin-GCaMP6 for PSD95, respectively), and Halo-tagged STIMs. Labelling intensity and duration as well as image acquisition rate was adjusted to gain single molecule resolution (see Methods, **Fig. 1A**). The dynamics of STIM proteins were assessed by SPT experiments at DIV 14-16. Coarse grain trajectories at a temporal resolution of 20 Hz were reconstructed and used to calculate the mean square displacement (MSD), diffusion coefficient (D) and local confinement of STIM proteins in the ER membrane. Overlap of trajectories with synaptic markers was used to assess synaptic localization of STIM proteins (**Fig. 1B, E**). At resting conditions using the overexpression approach, STIM1 exhibited significantly greater mobility in axons compared to dendrites (**Fig. 1C**). Store depletion by 20 µM cyclopiazonic acid (CPA) induced significant shifts in STIM1 diffusion in dendrites, however, no such changes were seen in axons (**Fig. 1D**). STIM2 protein showed more confined trajectories within dendrites under resting conditions (**Fig. 1E**) and similar to STIM1, STIM2 exhibited faster diffusion in axons than dendrites (**Fig. 1F**). Surprisingly, STIM2 dynamics remained unchanged in both compartments following CPA-induced store depletion (**Fig. 1G**). STIM2 is a more calcium-sensitive isoform and is observed to be in a pre-clustered state compared to STIM1, even without store depletion in non-excitable cells (Ong et al., 2015). This is reflected in the differences in the diffusion coefficients for dendritic STIM2 versus STIM1 (mean ± SD, STIM1 dendrite: D Avg 0.030 ± 0.004 µm^2^/s, STIM2 dendrite: D Avg 0.026 ± 0.006 µm^2^/s) (**Fig. 1H**).

**Figure 1.**
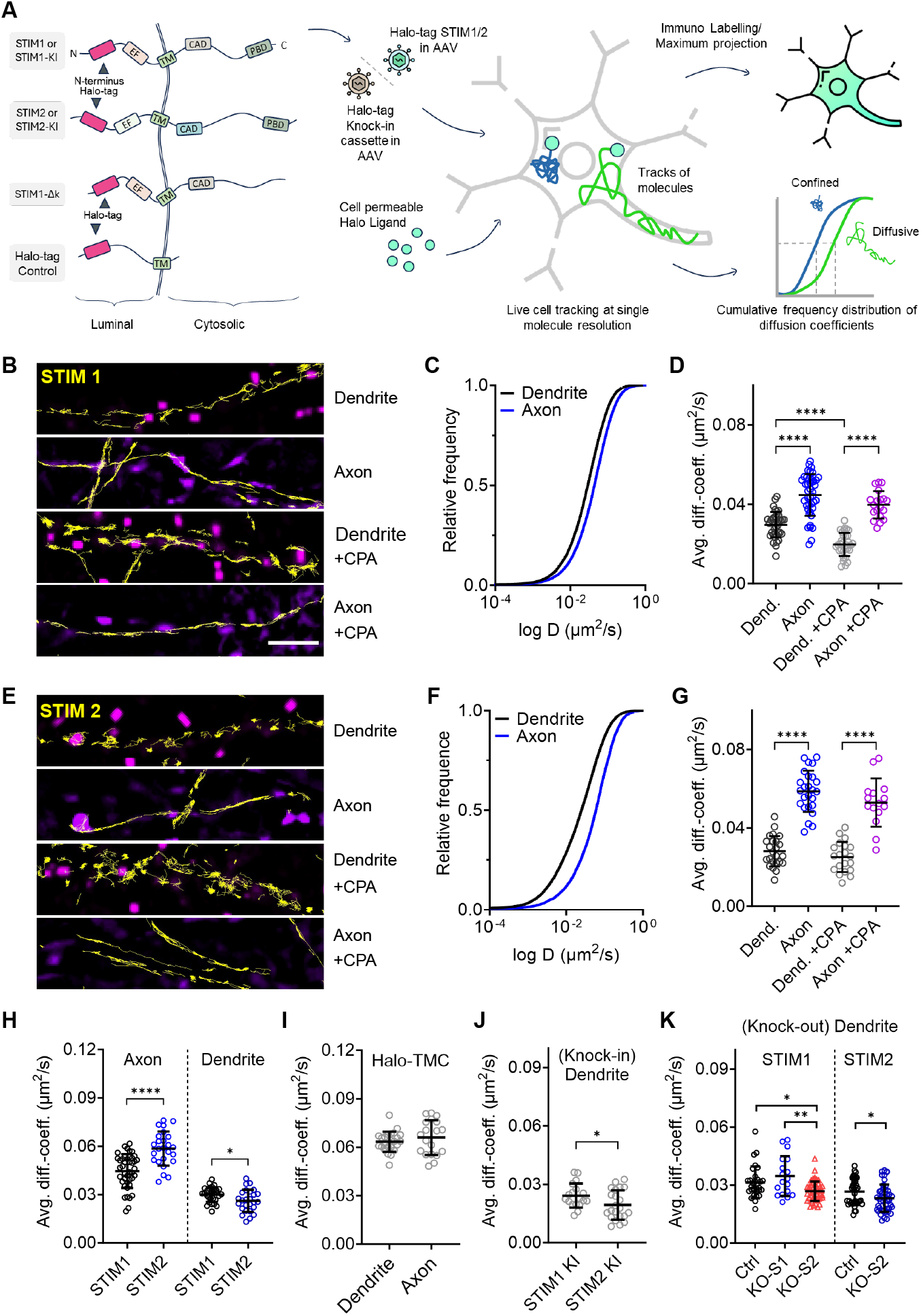
Differential dynamics of STIMs in neuronal compartments. **A)** Schematic representation of the constructs used in this study, highlighting the locations of tags and key domains of STIM proteins. The schematic also illustrates the AAV delivery approach and HaloTag labeling methodology employed for tracking the overexpressed or knock-in constructs. The diffusion coefficients derived from tracked molecules are plotted as cumulative frequency distributions on a logarithmic scale, illustrating the global relative changes in diffusion dynamics. **B)** Visualization of STIM1 tracking (yellow) within axons and dendrites, juxtaposed against presynaptic synaptophysin-GCaMP labeling (magenta in axon) and post-synaptic PSD95 labeling (magenta in dendrite). Data acquired for both resting conditions at 37°C (Ringer solution with 2 mM Ca^2+^, 2 mM Mg^2+^, 10 M CNQX, 10 M APV) and store-depleted conditions (Ringer solution with 2 mM Ca^2+^, 2 mM Mg^2+^, 10 M CNQX, 10 M APV, 20 M CPA). **C)** Cumulative frequency distribution showing the diffusion coefficient of STIM1 within axons and dendrites. **D)** Median diffusion coefficient comparison of STIM1 between the resting state and store-depleted conditions. Each data point depicted on the graph is the median diffusion coefficient from a single acquisition. **E)** Visual representation of Halo-STIM2 tracking (yellow) within axons and dendrites under both resting and store-depleted conditions, in relation to presynaptic synaptophysin-GCaMP and post-synaptic PSD95 labeling. **F)** Cumulative frequency distribution demonstrating the diffusion coefficient of STIM2 within axons and dendrites. **G)** Comparative evaluation of median diffusion coefficient for STIM2 between the resting state and store-depleted condition. Each data point presented on the graph represents the median from a single acquisition. **H)** Comparison of STIM1 and STIM2 diffusion within axons and dendrites, highlighting the presence of a fast-moving fraction in STIM2 in axons while a confined STIM2 in dendrites relative to STIM1. **I)** Diffusion of Halo-tag control (Halo-TMC) in axons and dendrites in resting conditions. **J)** SPT of STIM1 and STIM2 knock-in constructs, demonstrating a consistent trend of STIM2 being more confined than STIM1 in dendrites. **K)** Tracking of overexpressed STIM proteins was performed following Cre-mediated knockout of endogenous Stim1 and Stim2 (in the graphs, abbreviated as S1 and S2 respectively). The terms KO-S1 and KO-S2 refer to the Cre-mediated knockout of Stim1 and Stim2, respectively. rAAV carrying Cre was delivered to neurons on DIV2, while rAAV encoding Halo-STIM1 and Halo-STIM2 was introduced on DIV7. Imaging and data acquisition were performed on DIV14-15. Error bars for all figures depict the mean ± standard deviation (SD). Statistical significance is indicated as \*\*\*\**P* < 0.0001, \*\*\**P* < 0.001, \**P* < 0.1. For comparisons between two groups, a parametric unpaired t-test was used, while multiple groups were analyzed using one-way ANOVA followed by Tukey’s multiple comparisons test. Refer to the supplementary table for *p*-values, *N, n*, and trajectory counts.

However, in axons, STIM2 is more mobile (mean ± SD, STIM1 axon: D Avg 0.044 ± 0.01 µm^2^/s versus STIM2 axon: D Avg 0.058 ± 0.01 µm^2^/s) (**Fig. 1H**), suggesting that factors other than the local ER calcium concentration influence STIM mobility in axons and dendrites. Several possibilities could account for these differing diffusion properties including the variation in the tubular architecture of the ER in dendrites and axons (Y. Wu et al., 2017) which could potentially influence the diffusion properties of ER membrane-associated proteins (Domanov et al., 2011). To investigate this, we used a Halo-tagged control construct comprising both the STIM1 signal peptide and the ER transmembrane C-terminus of STIM1 (referred to as Halo-TMC) (**Fig. 1A**). This ER-membrane construct exhibited similar mobility in both axons and dendrites and did not form clusters in either compartment

Based on these findings, we conclude that structural differences in ER tubules diameter between dendrites and axons are unlikely to account for the observed differences in STIMs diffusion. Another reason for the drastically altered mobility of membrane-associated proteins could be an overexpression and the position of the tag in the constructs (Knight et al., 2010; Ramadurai et al., 2009). While tagging does not appear to affect the function of STIM proteins, as indicated by CPA-induced clustering (see above), overexpression can significantly increase surface mobility (Klatt et al., 2021; Lee et al., 2023). We performed two control experiments: first, examining the diffusion of overexpressed STIM proteins in a knockout (KO) background to eliminate the endogenous population, and second, tagging and tracking the endogenous STIM proteins. The dynamics of SITM1 and STIM2 remained similar after KO and re-expression of the respective isoform (**Fig. 1K**). However, STIM2 KO significantly reduced STIM1 dynamics (STIM1 dendrite Control: D Avg 0.031 ± 0.008 µm^2^/s, STIM1 dendrite in KO of STIM2: D Avg 0.026 ± 0.005 µm^2^/s, which is similar to STIM2 in resting conditions), suggesting STIM1 compensates for the absence of STIM2. This highlights isoform redundancy and functional role reversal, consistent with observations in T-cells (Oh-hora et al., 2008).

The endogenous STIM protein population was tagged using the CRISPR-Cas9 based tagging strategy ORANGE (Willems et al., 2020). Here we used a floxed Cas9-GFP mouse-line. Knock-ins were induced by dual rAAV infection of hippocampal neurons with one virus expressing the Cre-recombinase and another rAAV virus expressing the guide RNA for the specific insertion of the Halo-tag close to the N-terminal position used for the overexpression construct. Combining rAAV mediated expression of the Cre-recombinase and the STIM-specific guide RNA at 3-4 DIV led to a robust expression of tagged STIM proteins in a small fraction of neurons (corresponding to < 1% neurons per coverslip). In dendrites, the diffusion properties of labeled endogenous STIMs were consistent with those observed under overexpression conditions. STIM2 exhibited significantly lower diffusion than STIM1, confirming the validity of overexpression approach (**Fig. 1J**). However, tracking endogenous STIM proteins in axons was challenging due to their low abundance in these regions.

### 2.2. STIM protein are transient visitors in synaptic compartments

Previous studies using confocal imaging approaches showed partial colocalization of STIM proteins in pre- and postsynaptic compartments based on antibody-immunolabeling of endogenous STIM populations with significant variability (Chanaday et al., 2021; de Juan-Sanz et al., 2017; Kushnireva et al., 2021). Since the ER is notably absent from secretion-active zones (Y. Wu et al., 2017), we probed the abundance of STIMs by using both SPT and endogenous labeling (knock-ins) of STIMs. We initiated our investigation by analyzing STIM knock-ins using immunolabeling in combination with the presynaptic protein RIM1 (**Fig. 2A**). Evaluation of the Pearson’s correlation coefficient revealed a positive correlation between STIMs and RIM1, consistent with previous studies (**Fig. 2B**). However, the Manders’ overlap coefficients were notably low (mean ± SD: M1 STIM1: 0.09 ± 0.04, M1 STIM2: 0.18 ± 0.1, M2 STIM1: 0.04 ± 0.01, M2 STIM2: 0.07 ± 0.01, **Fig. 2C, D**), indicating that while the signals may be spatially adjacent, they are not truly colocalized. Considering potential artifacts associated with immunolabeling, these results imply that peri-synaptic regions might be misclassified as synaptic. Depletion of ER Ca^2+^ with CPA resulted in a modest increase in the Pearson’s coefficient for both STIM1 and STIM2. In contrast, the Manders’ overlap coefficients remained largely unchanged, which further prompted us to investigate the STIM-synaptic relationship in greater detail using SPT.

**Figure 2.**
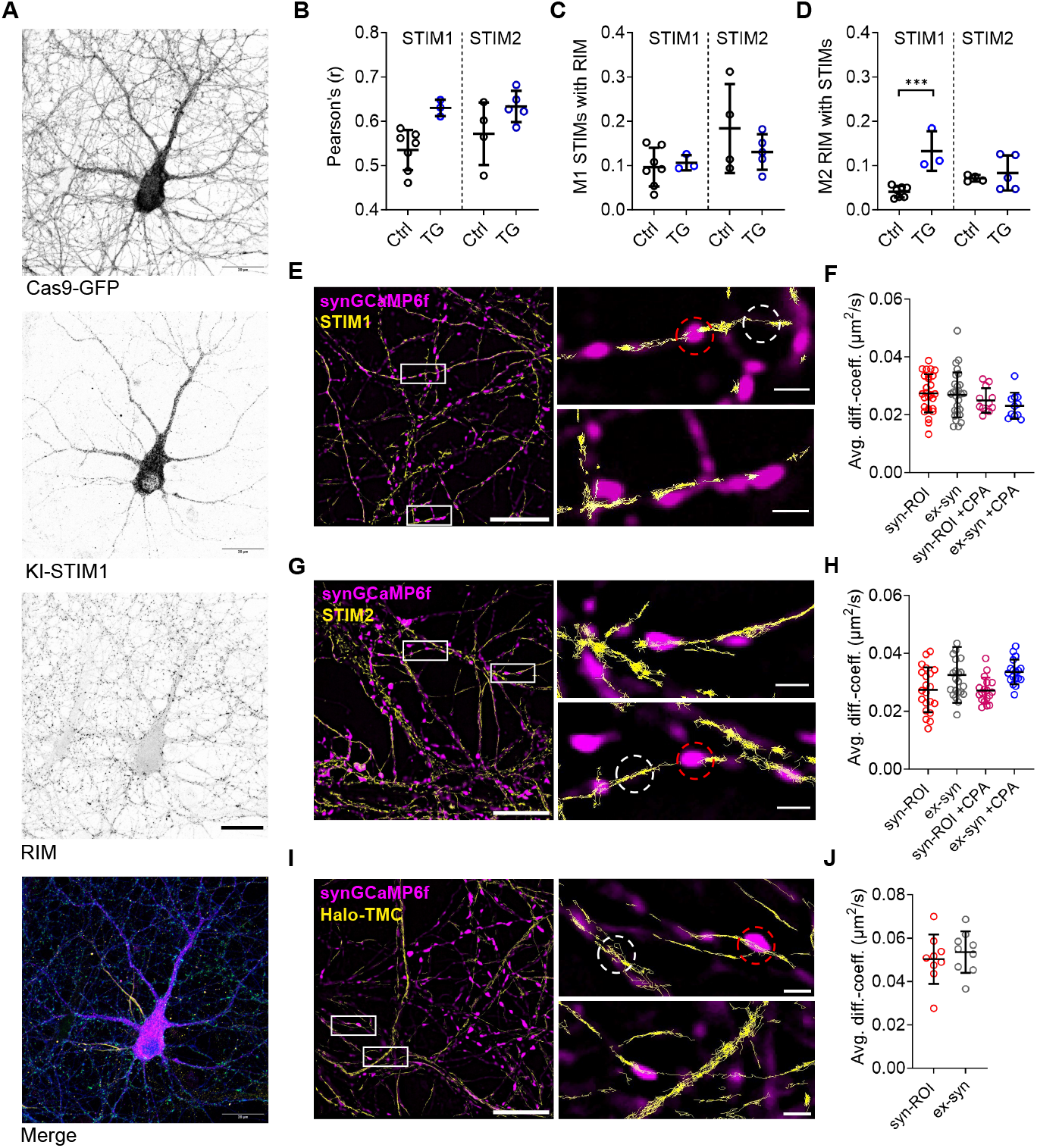
Presynaptic distribution of STIMs. **A)** Confocal imaging of knock-in Halo-STIM1 in Cas9-expressing hippocampal cultures, co-labeled with the presynaptic marker RIM. The merged image additionally displays AnkyrinG staining, not included in grayscale images and marked in yellow. Scale: 20 m. **B)** Pearson correlation coefficients for STIMs with RIM1. **C)** Manders’ M1 coefficients (co-localization of STIMs with RIM). **D)** Manders’ M2 coefficients (co-localization of RIM with STIMs) calculated for both STIM1 and STIM2 under resting and store-depleted conditions (1 M TG). **E, G, I)** Visualization of STIM1 (**E**), STIM2 (**G**), and Halo-control (**I**) tracks (yellow) in axons co-labeled with presynaptic synaptophysin-GCaMP. Representative 0.91 m^2^ regions of interest (ROIs) are marked as synaptic (red) and extrasynaptic (white). Scale: 10 m for full images, 2 m for magnified views. **F, H, J)** Median diffusion coefficients of STIM1 (**F**), STIM2 (**H**), and Halo-control (**J**) plotted under resting conditions at 37°C (Ringer’s solution with 2 mM Ca^2+^, 2 mM Mg^2+^, 10 M CNQX, 10 M APV) and store-depleted conditions (Ringer’s solution with 20 M CPA). Each data point represents the median diffusion coefficient from a single acquisition encompassing multiple presynaptic locations. Error bars indicate mean ± SD. Refer to the supplementary table for *p*-values, *N, n*, and trajectory counts.

We labeled the presynaptic compartment with synaptophysin-GCaMP6 delivered via rAAV, alongside either STIM1 or STIM2. Interestingly, STIM protein tracks did not accumulate within the core of the presynaptic compartments. However, STIMs occasionally passed through these regions, potentially due to the continuous network of ER tubules (**Fig. 2E, G**). For these infrequent events, we quantified changes in diffusion following store depletion. For analysis, regions of interest (ROIs) with a 15-pixel diameter (1.08 m, corresponding to an area of 0.91 m^2^) were defined around synaptophysin-GCaMP6 spots, as shown in the enlarged section of **Fig. 2E, G, I**. These ROIs encompassed not only the synaptic core but also the adjacent peri-synaptic regions, referred to as syn-ROI in the graphs. In comparison, regions outside the synaptic area were analyzed separately and are termed extra-synaptic (or ex-syn in the graphs). Store depletion with CPA did not alter the diffusion properties of either STIM1 or STIM2 in synaptic and extra-synaptic ROIs, suggesting that ER calcium content does not promote the recruitment of STIMs into presynaptic active zones (**Fig. 2F, H**).

Tracking the movement of STIM1 and STIM2 showed that both proteins were not confined within the presynaptic core but frequently localized to peri-synaptic regions. In contrast, the Halo-TMC exhibited uniform trajectories and diffusion both in and outside synapses, with no significant enrichment of confined tracks in the peri-synaptic region (**Fig. 2I, J**). This pattern suggests that STIMs preferentially localize to peri-synaptic sites, likely due to the structural arrangement of the ER near the synapse, which may facilitate interactions between STIMs and plasma membrane proteins in this region.

In the postsynaptic compartment, events of STIM proteins localizing in close proximity to PSD-95 were extremely rare, with no evident enrichment in the PSD. Maximum projections of SPT movies revealed smooth, tube-like structures for both STIM1 and STIM2. Notably, STIM1 displayed a uniform distribution, while STIM2 exhibited a punctate pattern in dendrites (**Fig. 3A, B**). To assess whether STIM proteins avoid entering ER structures like the spine apparatus or if synaptic ER extensions are absent in our cultured neurons, we used a Halo-TMC as a control, which clearly outlined the ER, frequently highlighting the spine apparatus and occasionally showing ER structures near PSD-95 spots (**Fig. 3C**). Quantitative analysis indicated that ER visits, visualized by the Halo-TMC, occurred in 50% of spines, whereas STIM proteins were rarely detected (<5%) (**Sup. Fig. 3**).

**Figure 3.**
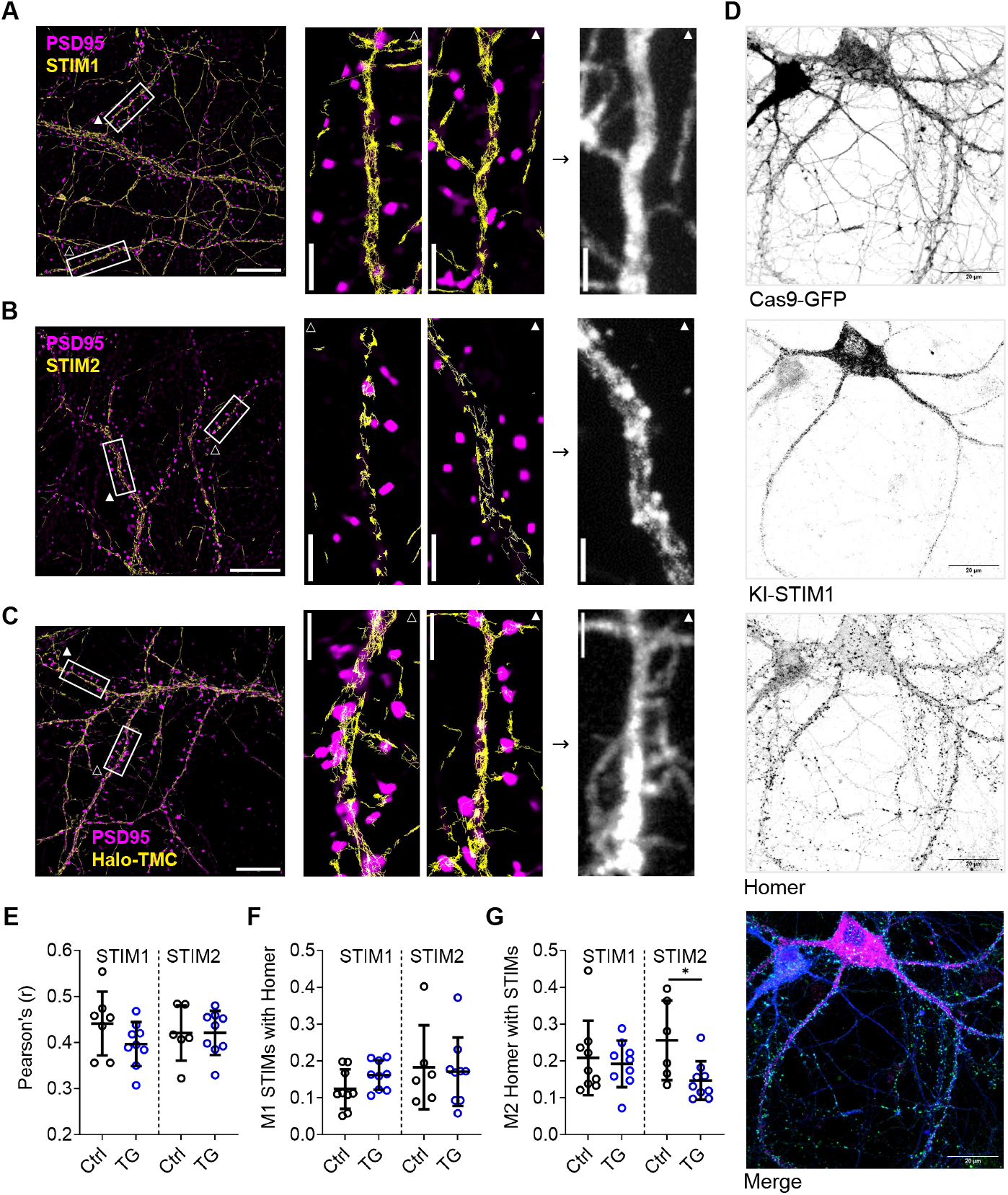
Post-synaptic distribution of STIMs. **A-C)** Visualization of Halo-STIM1 (**A**), STIM2 (**B**), and Halo-control (**C**) tracks (yellow) in dendrites co-labeled with the postsynaptic marker PSD-95. Acquisitions were performed under resting conditions at 37 °C (Ringer’s solution with 2 mM Ca^2+^, 2 mM Mg^2+^, 10 M CNQX, 10 M APV). The grayscale images represent maximum projections of SPT movies without synaptic labels in magnified dendritic sections. Scale bars: 10 m (large images), 2 m (magnified views). **D)** Confocal imaging of knock-in Halo-STIM1 in Cas9-expressing hippocampal cultures, co-labeled with the post-synaptic marker Homer. Scale 20 m. **E)** Pearson correlation coefficients of STIMs with Homer, **F)** Manders’ *M*_1_ coefficients (co-localization of STIMs with Homer), and **G)** Manders’ *M*_2_ coefficients (co-localization of Homer with STIMs) calculated for both STIM1 and STIM2 under resting and store-depleted conditions (TG). Error bars in graphs indicate the mean ± SD. Refer to supplementary table for p-values, *N, n*, and trajectory counts.

Fixed immunostaining of endogenously labeled STIM proteins alongside the postsynaptic marker Homer (**Fig. 3D**) confirmed our SPT observations. Although the Pearson’s coefficient revealed a weak positive correlation between Homer and STIM signals, the values of Manders’ overlap coefficients were very low (**Fig. 3E**). Both Pearson’s and Manders coefficients remained unchanged after store depletion with TG (thapsigargin), suggesting that STIM proteins are not recruited to the postsynaptic spine upon ER Ca^2+^ store depletion (**Fig. 3F-G**). Collectively, the results from both knock-in immunostaining and SPT suggest that STIM proteins are not stabilized within synapses and are much more frequent in clusters along the dendritic shaft, posing the question of whether STIM proteins contribute directly to synaptic signaling.

### 2.3. Chronic and acute changes in neuronal activity alter STIM dynamics in dendrites

While STIM proteins are not strictly localized to synaptic compartments, it is reasonable to hypothesize that synaptic activity influences ER-PM communication and thereby modulates STIM protein dynamics. Emerging evidence indicates that STIM proteins and the ER play a critical role in homeostatic plasticity through the regulation of calcium signaling (Chanaday & Kavalali, 2022; Maggio & Vlachos, 2014). In our SPT experiments, neurons subjected to 24-hour Tetrodotoxin (TTX) or Bicuculline (BIC) treatments showed that altered network activity significantly influenced STIM1 mobility within dendrites: high activity (BIC) restricted STIM1 movement, whereas low activity (TTX) increased it. Interestingly, STIM1 in axons and STIM2 in both compartments remained unaffected by activity changes (**Sup Fig. 4**), highlighting distinct activity-dependent behavior for STIM1 and pre-formed stability for STIM2. Additionally, the TTX-induced effects suggest the involvement of activity-regulating proteins in modulating STIM1 function. To further investigate this, we applied glutamate for acute treatments, leveraging its well-established role in modulating neuronal activity and its documented connection to STIM proteins (Dittmer et al., 2017; Dittmer & Dell’Acqua, 2024).

We hypothesized that calcium influx through NMDA receptors, which triggers calcium release from the ER via CICR, might be crucial for this process. We observed that brief (1 min) exposure to 20 µM glutamate caused a robust clustering of dendritic STIM1. Notably, blocking NMDA receptors using APV alone was sufficient to prevent the effect of glutamate on STIM dynamics (**Fig. 4A-C**). Maximum projections of SPT data of STIM1 showed a uniform signal across the dendrite in the control condition or during blockade of NMDAR activation by APV. Diffusion coefficients of STIM1 were similar under NMDA blockade or control conditions (**Fig. 4B**). In contrast, glutamate treatment without NMDA receptor blockade induced significant clustering of STIM1 in dendrites (**Fig. 4A-C**). To ensure that only NMDA receptors were involved in the clustering phenomenon, the potential contribution of other glutamate receptors, namely AMPA and group I metabotropic glutamate receptors, was also examined. Activation of these receptors using specific agonists, AMPA and DHPG respectively, did not induce any changes in STIM1 dynamics in dendrites (**Sup. Fig. 4**), which contrasts with earlier FRAP experiments (Ng, Krogh, and Toresson, 2011).

**Figure 4.**
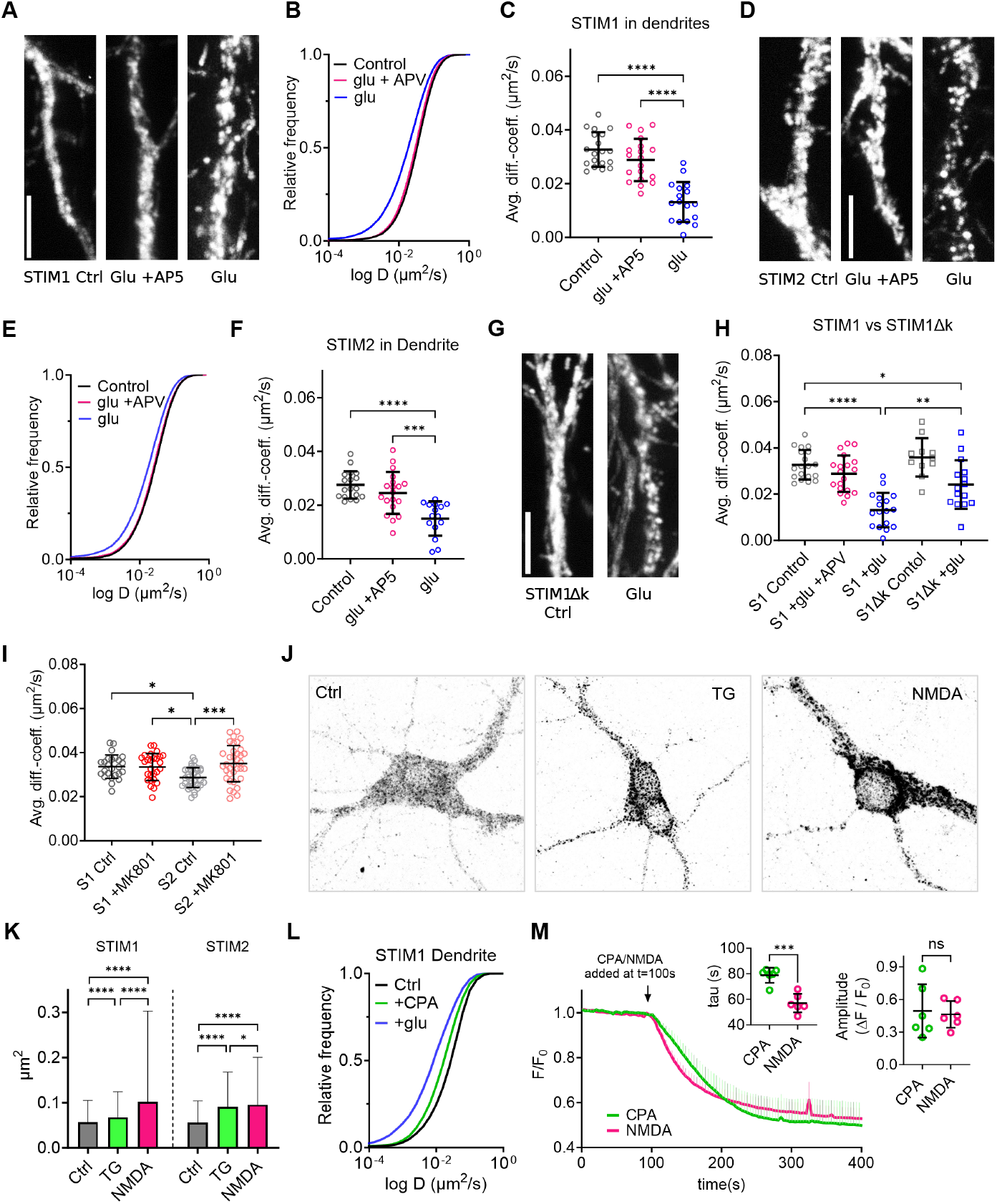
Glutamate-induced clustering of STIM proteins depends on NMDA signaling and neuronal activity. Neurons were exposed to 20 M glutamate (Glu) in Mg^2+^-free Ringer solution prior to imaging, followed by wash and acquisition in Ringer solution (including Mg^2+^) supplemented with 10 M APV and 10 M CNQX at 37 ^°^C. Representative maximum projection images of 500 frames extracted from STIM1 (**A**), STIM2 (**D**), and STIM1K (**G**) SPT movies showing dendrites under control conditions, glutamate treatment, and NMDA receptor blockade with APV. **B, E)** Cumulative frequency distribution plot of STIM1 (**B**) and STIM2 (**E**) diffusion coefficients (*N* = 3 for both). **C, F)** Comparative evaluation of median diffusion coefficients for STIM1 (**C**) and STIM2 (**F**) across conditions, with each data point representing the median from a single acquisition. **G)** Maximum projection images of STIM1K under control conditions and after treatment with glutamate. **H)** Median diffusion coefficients of STIM1K, including comparisons with the diffusion data for wild-type STIM1. **I)** Statistical analysis and comparison of diffusion coefficients of both STIM1 and STIM2 after 24 h MK-801 treatment. **J)** Immunostaining of endogenously labeled STIM1 in Cas9 neurons, comparing STIM1 cluster morphology under store depletion with TG treatment and NMDA treatment. **K)** Cluster size quantification of STIM knock-ins under Control, CPA, and NMDA treatments, analyzed from 4–5 cells per condition. **L)** SPT analysis of STIM1, comparing the impact of glutamate on STIM1 dynamics. Cumulative frequency distribution plot showing the diffusion coefficients of STIM1 during treatments with glutamate (20 M) and CPA (20 M). **M)** ER-GCaMP Ca^2+^ imaging in hippocampal neurons treated with CPA or NMDA to induce Ca^2+^ release from the ER at *t* = 100 s. The figure compares the amplitude of ER Ca^2+^ release and the kinetics of Ca^2+^ release (*τ*) between the two treatments. Error bars for all figures depict the mean ± standard deviation. Statistical significance is indicated as \*\*\*\**P* < 0.0001, \*\*\**P* < 0.001, \**P* < 0.1. For comparisons between two groups, a parametric unpaired *t*-test was used, while multiple groups were analyzed using one-way ANOVA followed by Tukey’s multiple comparisons test. Refer to supplementary table for *p*-values, *N, n*, and trajectory counts.

Similar observations were made for STIM2 dynamics in dendrites. Blocking NMDA receptors with APV prevented glutamate-induced STIM2 clustering in dendrites (**Fig. 4D-F**). The magnitude of STIM1 diffusion changes appeared greater than that of STIM2, signifying the broader dynamic range of STIM1 compared to STIM2. Further, we investigated whether glutamate-mediated clustering of STIM proteins is facilitated by tighter association with the PM. To explore this, we utilized STIM1Δk, a mutant lacking the PM-binding PDB domain of STIM1. Upon glutamate treatment, the clustering of STIM1Δk was less robust compared to wild-type STIM1 (**Fig. 4G, H**), indicating that the translocation of STIM proteins to ER-PM junctions contributes to clustering but is not the sole significant factor.

The differences in the confinement of STIM1 and STIM2 under control conditions (**Fig. 1H-K**) prompted us to investigate whether spontaneous activity of NMDARs might already induce clustering of STIM2 due to the difference in calcium affinity of the two isoforms. Overnight application of MK-801, an irreversible NMDAR antagonist, resulted in a more diffuse distribution of STIM2 (**Fig. 4I**), indicating a role for basal NMDAR activity in the clustering of STIM2. In contrast, no changes were observed in axonal STIM1 or STIM2 dynamics in response to glutamate treatment (**Sup Fig. 4**).

Acute glutamate treatment resulted in significantly stronger clustering of STIM proteins compared to store depletion induced by CPA (**Fig. 4L**). Additionally, the morphology of STIM clusters in knock-ins triggered by NMDA differed significantly from those formed by passive calcium leak from the ER due to TG-induced SERCA inactivation. Both STIM1 and STIM2 cluster areas were significantly larger following NMDA treatment compared to TG treatment (**Fig. 4J, K**). Ryanodine-induced depletion of ER stores via activation of the ryanodine receptor (RyR) resulted in less pronounced STIM clustering compared to NMDA or glutamate treatment (**Sup. Fig. 8**). According to the established model of STIM function, the degree of store depletion, irrespective of the depletion method, should correlate with the extent of STIM clustering and activation (Zhou et al., 2013). To validate this, we compared store depletion induced by NMDA and CPA using ER-GCaMP Ca^2^ imaging. Both treatments resulted in similar final depletion levels (**Fig. 4M**). However, the rate of ER Ca^2^ depletion was faster upon NMDA treatment compared with CPA. This suggests that the dynamics of ER Ca^2^ store depletion, and possibly other factors, determine the clustering of STIM molecules in neurons. In the next section, we further explore the mechanisms of activation of neuronal STIMs.

### 2.4. Glutamate-induced STIM clustering is not dependent on the activity of Cav1.2

Apart from the classical STIM-Orai interaction, several studies suggest a functional relationship between STIMs and Ca_v_1.2. Both STIM isoforms have been shown to directly inhibit Ca_v_1.2 activity (Park et al., 2010; Wang et al., 2010). Direct glutamate uncaging experiments reveal that STIM proteins, particularly STIM1, play a key role in regulating Ca_v_1.2 channel activity (Dittmer et al., 2017). Through direct somatic calcium current measurements, local dendritic calcium imaging, and shRNAi-mediated suppression of STIM1, the following signaling pathway has been proposed: NMDA receptor activation, along with AMPAR-induced depolarization, opens synaptic and peri-synaptic Ca_v_1.2 channels which subsequently trigger CICR and STIM1 clustering. STIM1 transiently clusters following glutamate application and inhibits Ca_v_1.2 channels, thereby preventing excessive Ca^2+^ influx into the postsynaptic compartment. Notably, direct Ca_v_1.2 inhibition was shown to reduce STIM1 clustering (Dittmer et al., 2017).

We hypothesized that blocking Ca_v_1.2 channels would result in a noticeable difference in glutamate-induced STIM protein clustering. To test this, we tracked STIM dynamics following NMDA treatment in the presence of the Ca_v_1.2 channel blocker nimodipine (nim). However, nimodipine did not inhibit STIM1 cluster formation (**Fig. 5A**). Additionally, activation of Ca_v_1.2 channels with BayK also had no detectable effect on STIM protein mobility (**Sup. Fig. 4C**). Furthermore, analysis of STIM1 mobility revealed no significant difference between nimodipine-pretreated neurons and control neurons (**Fig. 5B**). This suggests that NMDA-mediated STIM1 clustering occurs independently of LTCCs.

**Figure 5.**
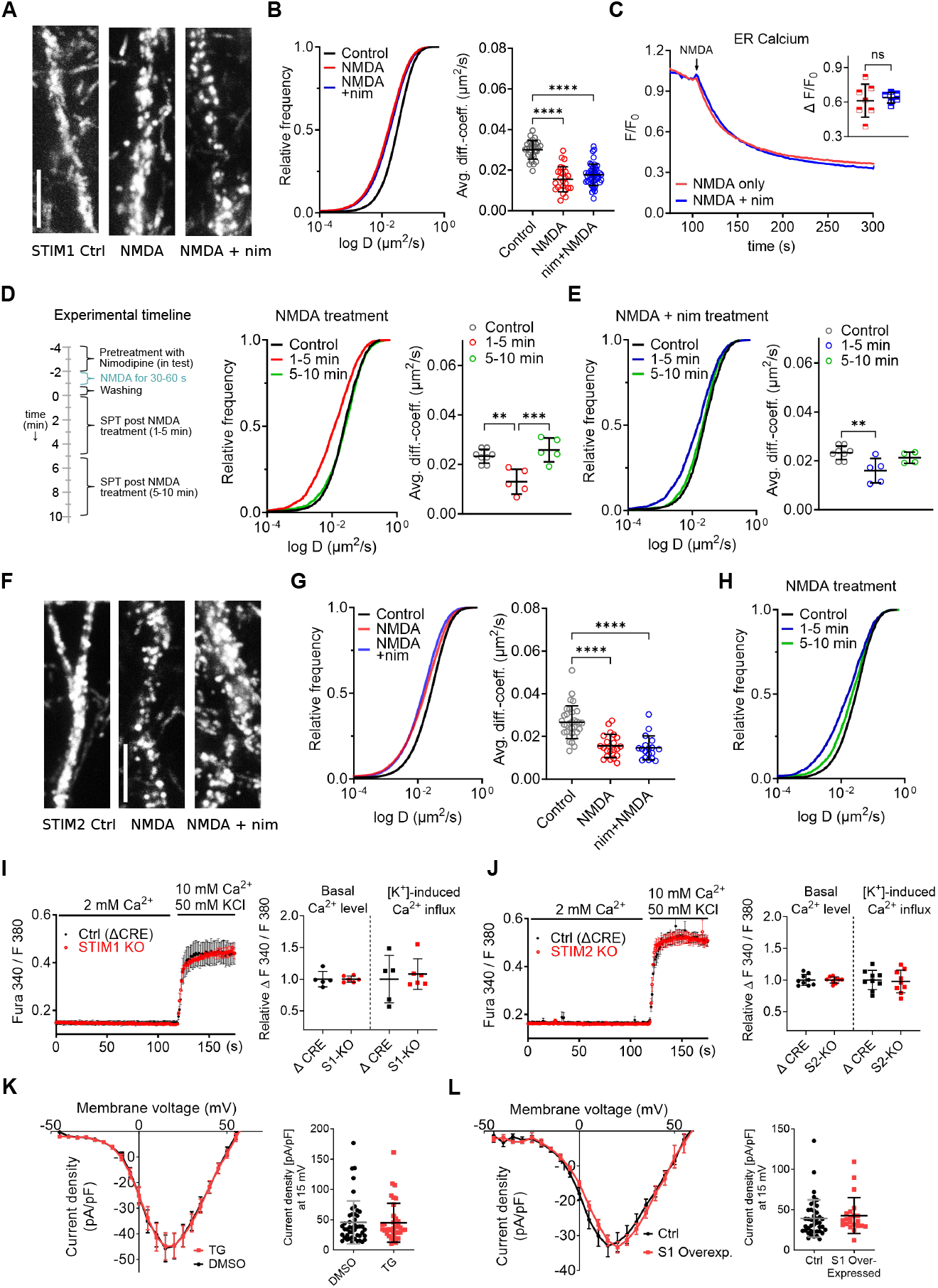
Contribution of L-type calcium channels in NMDA induced clustering of STIMs. **A)** Maximum intensity projection images from STIM1 SPT movies under control conditions, NMDA treatment, and NMDA + nimodipine treatment. NMDA stimulation involved 50 M NMDA in Mg^2+^- and APV-free Ringer solution for 1 minute, followed by washout. For nimodipine treatment, cells were pre-incubated with 50 M nimodipine (nim) for 3 minutes prior to and during NMDA activation. Imaging was performed in Ringer solution containing 2 mM Mg^2+^, 2 mM Ca^2+^, and 10 M CNQX at 37 ^°^C. **B)** Cumulative frequency distribution plot and statistical analysis of median diffusion coefficients per acquisition for STIM1. **C)** ER calcium imaging using ER-GCaMP-150, with NMDA added at *t* = 100 s for both control and nimodipine pre-treated conditions (see Methods). **D–E)** Experimental timeline for assessing the temporal dynamics of STIMs following NMDA treatment. **D)** Cumulative distribution plot of diffusion coefficients and median diffusion coefficients per acquisition for STIM1 under NMDA treatments. **E)** Corresponding data for the nimodipine pre-treatment condition. Control recordings primarily included data from the first 1–5 minutes of acquisition. For NMDA- and nimodipine + NMDA-treated cells, continuous recordings were performed for 10 minutes, with data separated and plotted as two distinct time intervals: 1–5 minutes and 5–10 minutes post-NMDA treatment, collected from the same coverslip. **F)** Representative maximum projection images from STIM2 SPT movies under the same treatments and conditions as described for STIM1. **G)** Cumulative frequency distribution plot and statistical analysis of median diffusion coefficients per acquisition for STIM2. **H)** Temporal dynamics of STIM2 following NMDA treatment, with data separated and plotted as two distinct time intervals: 1–5 minutes and 5–10 minutes post-NMDA treatment, obtained from the same coverslip. **I**,**J)** Cytosolic calcium levels were measured using Fura-2 in hippocampal neurons with Stim1 (**I**) and Stim2 (**J**) knockouts. Experiments were performed at 37 ^°^C in Ringer’s solution, with a high-potassium pulse introduced at *t* = 120 s. Data were collected from neurons at DIV 14–16. **K**,**L)** Whole-cell patch-clamp recordings from HEK293T cells stably expressing L-type 1.2 calcium channels (see Methods). **K)** Measurement of calcium currents following store depletion induced by 1 M TG or with vehicle (DMSO). **L)** Measurement of calcium currents in cells overexpressing EYFP-STIM1 in comparison to control. Refer to supplementary table for *p*-values, *N, n*, and trajectory counts.

We further investigated the effect of LTCC blockade on ER Ca^2+^. Nimodipine did not alter the kinetics or the magnitude of ER Ca^2+^ efflux triggered by NMDA (**Fig. 5C**), suggesting no major impact of Ca_v_1.2 channel activity on the NMDAR-induced changes in ER-calcium. To evaluate the specificity of the NMDAR-induced STIM1 clustering further, we explored the duration of STIM clustering after washout of NMDA. NMDA triggered STIM1 clustering persisted for approximately 5 minutes. Subsequently, the dynamics of STIM1 returned to a basal level within the next 10 minutes (**Fig. 5D**). This timeframe (5–10 minutes) aligns with reported effects of glutamate on ER calcium levels (Dittmer et al., 2017). We replicated the time course in nimodipine-pretreated neurons followed by NMDA exposure. Similar to the NMDA-only treatment, STIM1 clustered for 5 minutes before reverting to normal diffusion by 10 minutes (**Fig. 5E**). These experiments confirm the transient nature of STIM1 clustering, as reported by Dittmer et al. (2017), and further emphasize that STIM1 clustering is independent of Ca_v_1.2 channel activity.

Similar results were obtained for STIM2. Nimodipine did not prevent the robust clustering of STIM2 triggered by NMDA treatment, which, like STIM1, was observed within seconds and is transient (**Fig. 5F–H**). The maximum projection images revealed a clear shift in STIM2 cluster formation upon NMDA exposure, and this clustering pattern remained unchanged when LTCCs were inhibited alongside NMDA activation (**Fig. 5F**). Furthermore, analysis of STIM2 dynamics under both conditions showed no significant differences, confirming that nimodipine had no effect on NMDA-mediated clustering of either STIM1 or STIM2 (**Fig. 5G**). This suggests that LTCCs are not crucial for NMDA-induced clustering of either STIM1 or STIM2.

The persistent clustering of STIM2 proteins under control conditions could indicate that a fraction of ER-PM junctions composed of STIM2-ORAI may exist within neurons to support replenishment of ongoing calcium fluxes during neuronal network activity. We accessed this by use of Fura-2 imaging to measure cytosolic calcium dynamics in control and STIM knockout neurons. Neurons were infected with either an rAAV Cre virus or a control virus expressing an inactive, mutated Cre (Cre), both carrying a bicistronic TdTomato reporter signal (**Sup. Fig. 5**). Knockout of either STIM1 or STIM2 did not affect basal cytosolic Ca^2+^ concentration, which suggests that SOCE is not a critical determinant of resting cytoplasmic [Ca^2+^] in neurons (**Fig. 5I, J**).

Next, we depolarized the neuronal membrane by addition of KCl (50 mM). There was no impact on the induced calcium transients in both STIM1 and STIM2 KOs (**Fig. 5I, J**), speaking against the reported inhibition of Ca_v_1.2 channels by STIM proteins. The reduction in STIM-mediated inhibition of Ca_v_1.2 channels should result in a higher amplitude of depolarization-induced Ca^2+^ transients. However, KO of STIM1 or STIM2 had no influence on this parameter. Having determined that STIM proteins likely have no impact on the activity of Ca_v_1.2 in mouse hippocampal neurons, we directly investigated the potential negative regulation of Ca_v_1.2 by STIM1 using patch-clamp recordings on HEK cells stably expressing Ca_v_1.2. We confirmed that neither activation of endogenous STIM1 by ER Ca^2+^ store depletion nor overexpression of STIM1 altered the L-type currents (**Fig. 5K, L**). Altogether, the SPT, Ca^2+^ imaging of cytosolic and ER signals and patch-clamp data argue against a direct coupling of STIMs with Ca_v_1.2.

However, the tendency of STIM2 to cluster along dendrites, similar to the voltage-gated K_v_+ channel (K_v_2.1), which organizes signaling complexes containing LTCCs at the soma-dendritic ER–PM contacts, suggests a potential role of STIM2 in organizing similar signaling microdomains (Mandikian et al., 2014; Vierra et al., 2019, 2021). Given the unconfirmed functional interaction between VGCCs and STIMs in our experiments, these STIM2-positive ER-PM junctions may constitute a type of neuronal ER-PM contact that is distinct from K_v_2.1 dependent cluster. To explore this further, we investigated the colocalization of STIM proteins with K_v_2.1.

### 2.5. STIMs are not confined to one type of ER-PM contact

The persistent clustering of STIM2 proteins under control conditions could indicate that a fraction of ER-PM junctions composed of STIM2-ORAI may exist within neurons to support replenishment of ongoing calcium fluxes during neuronal network activity. We accessed this by use of Fura-2 imaging to measure cytosolic calcium dynamics in control and STIM knockout neurons. Neurons were infected with either an rAAV Cre virus or a control virus expressing an inactive, mutated Cre (Cre), both carrying a bicistronic TdTomato reporter signal (**Sup. Fig. 5**). Knockout of either STIM1 or STIM2 did not affect basal cytosolic Ca^2+^ concentration, which suggests that SOCE is not a critical determinant of resting cytoplasmic [Ca^2+^] in neurons (**Fig. 5I, J**).

Next, we depolarized the neuronal membrane by addition of KCl (50 mM). There was no impact on the induced calcium transients in both STIM1 and STIM2 KOs (**Fig. 5I, J**), speaking against the reported inhibition of Ca_v_1.2 channels by STIM proteins. The reduction in STIM-mediated inhibition of Ca_v_1.2 channels should result in a higher amplitude of depolarization-induced Ca^2+^ transients. However, KO of STIM1 or STIM2 had no influence on this parameter. Having determined that STIM proteins likely have no impact on the activity of Ca_v_1.2 in mouse hippocampal neurons, we directly investigated the potential negative regulation of Ca_v_1.2 by STIM1 using patch-clamp recordings on HEK cells stably expressing Ca_v_1.2. We confirmed that neither activation of endogenous STIM1 by ER Ca^2+^ store depletion nor overexpression of STIM1 altered the L-type currents (**Fig. 5K, L**). Altogether, the SPT, Ca^2+^ imaging of cytosolic and ER signals and patch-clamp data argue against a direct coupling of STIMs with Ca_v_1.2.

However, the tendency of STIM2 to cluster along dendrites, similar to the voltage-gated K_v_+ channel (K_v_2.1), which organizes signaling complexes containing LTCCs at the soma-dendritic ER–PM contacts, suggests a potential role of STIM2 in organizing similar signaling microdomains (Mandikian et al., 2014; Vierra et al., 2019, 2021). Given the unconfirmed functional interaction between VGCCs and STIMs in our experiments, these STIM2-positive ER-PM junctions may constitute a type of neuronal ER-PM contact that is distinct from K_v_2.1 dependent cluster. To explore this further, we investigated the colocalization of STIM proteins with K_v_2.1. localization patterns.

## 3. Discussion

The precision of intracellular calcium signaling is a key element for many neuronal functions and has been particularly well-studied within chemical synapses between neurons. Although activity-evoked calcium signals in neurons have been extensively investigated, the relationship between the position and function of proteins assigned to SOCE in neurons remains obscure. Here, we employed SPT to examine the organization of STIM proteins within the ER membrane, as well as the timing and location of their connections to the plasma membrane. We leveraged the SPT approach to achieve the spatial and temporal resolution required to investigate synaptic compartments in detail, enabling a dynamic characterization of STIM protein behavior rather than relying on static snapshots taken before and after treatment. The well-established mobile organization of STIM proteins suggests only transient confinement of these proteins to spatially defined signaling compartments (Deng et al., 2009; Sallinger et al., 2023; Soboloff et al., 2012). Using three distinct approaches—monitoring endogenous, overexpressed, and re-expressed Halo-tagged STIM molecules—we demonstrate that STIM proteins are highly dynamic within the ER membrane and are transiently confined to ER-PM junctions.

The inherent differences in the sensitivity of STIM proteins to changes in ER luminal calcium concentration are reflected in the dynamics of the isoforms tested here, STIM1 and STIM2. Using a CRISPR-Cas9-based endogenous labeling strategy, we can rule out that the observed dynamics are biased by the use of specific splice variants or overexpression of STIM proteins (**Fig. 1**). Modulating neuronal network activity confirmed that STIM protein mobility is sensitive to neuronal excitability (**Fig. 4**) and is specifically fine-tuned by the tonic activity of NMDARs (**Fig. 5**). Strong activation of glutamatergic transmission directly links STIM protein mobility to fluctuations in ER calcium concentration. In this context, NMDA receptor activation serves as the predominant signal for confining STIM proteins to ER-PM junctions, as previously reported (Dittmer et al., 2017; Dittmer & Dell’Acqua, 2024). However, we did not find evidence to support the hypothesis that voltage-gated calcium channels, particularly CaV1.2 channels, significantly influence the confinement of STIM proteins.

The abundance of ER-PM junctions containing STIM proteins in close proximity to synapses may serve as a structural element to ensure that local calcium signals, triggered by the opening of NMDA receptors, are effectively relayed to ER effectors such as ryanodine receptors. It is reasonable to assume that STIM-mediated ER-PM contacts are both necessary and sufficient to link calcium signals between the plasma membrane and the ER, as extensively studied in other cell types (Ahmad, Narayanasamy, et al., 2022; Carrasco & Meyer, 2011). Since we found limited presence of STIM protein clusters within synapses, we propose that STIM-mediated peri-synaptic ER-PM junctions are adequate to modulate the coupling of synaptic calcium signals to ER calcium stores. This hypothesis aligns with findings from functional studies (Chanaday et al., 2021; de Juan-Sanz et al., 2017).

In HEK cells, clusters of immobile STIM2 were shown to act as sites of pre-defined Ca^2+^ signaling hubs that recruit STIM1 upon stimulation (Ahmad, Narayanasamy, et al., 2022). Our data suggest that a similar mechanism takes place in neurons. Indeed, using MAPPER, a genetically engineered marker of ER-PM junctions, we show that in unstimulated neurons, STIM2 clusters in MAPPER-positive spots, which means that clusters of STIM2 proteins represent ER-PM junctions. STIM1, on the other hand, is recruited to these pre-formed clusters only upon stronger ER Ca^2+^ store depletion (**Fig. 6**).Notably, neuronal stimulation did not affect the dynamics of axonal STIM molecules, suggesting heterogeneity of STIM functions in different neuronal compartments. It may therefore be hypothesized that STIM proteins serve, at least partially, differential roles in excitable and non-excitable cells. The importance of SOCE for non-excitable cells (reviewed in Rosado et al., 2016) and the controversy regarding the physiological relevance of SOCE in neurons (Lu & Fivaz, 2016) support this hypothesis. Indeed, although activation of ISOC and ICRAC currents is considered as a bona-fide function of STIM molecules, accumulating evidence suggests SOCE-independent roles of STIM molecules (Berna-Erro et al., 2009; de Juan-Sanz et al., 2017; Dittmer et al., 2017; Garcia-Alvarez, Shetty, et al., 2015; Lefkimmiatis et al., 2009; Majewski et al., 2017; Park et al., 2010; Tian et al., 2012; Wang et al., 2010). Similarly, although a number of synaptic phenotypes for STIM molecules have been proposed (Chanaday et al., 2021; de Juan-Sanz et al., 2017; Dittmer et al., 2017; Ramesh et al., 2021; S. Sun et al., 2014), direct evidence for the presence of ORAI proteins in boutons or spines remains sparse (Basnayake et al., 2021; Korkotian et al., 2017). It is not clear why the neuron would need such a low-conductance channel as ORAI to modulate neurotransmitter release or amplify the postsynaptic Ca^2+^ signal. There are indications for local replenishment of ER-compartments by SOCE mediated coupling within synaptic spines (S. Sun et al., 2014), however, our data do not strongly support this hypothesis, since we do not often observe confinement of STIM proteins in spines (**Fig. 3**). Indeed, our SPT data failed to reveal any enrichment of STIM proteins in the pre- or postsynaptic regions (**Fig. 2 E-J, Fig. 3 A-C, Sup. Fig. 3**). In fact, STIM molecules tended to avoid both the pre- and postsynaptic compartments and ER Ca^2+^ depletion did not affect the mobility of STIM molecules within the synapse. This is consistent with an electron-microscopy (EM) study, which showed that the ER does not come into direct contact with the active zone of neurotransmitter secretion (Y. Wu et al., 2017). Similarly, although the ER is thought to be present in around 40% of dendritic spines (Perez-Alvarez et al., 2020), EM data showed that the ER does not enter the PSD (Y. Wu et al., 2017). Our immunocytochemical experiments showed that colocalization between synaptic markers and endogenously halo-tagged STIMs was low (**Fig. 2 A-D**). STIM1 colocalization with the presynaptic marker RIM 1/2 did moderately increase upon ER store-depletion, nonetheless, Mander’s coefficient values suggest that STIMs are present only in a small subset of synapses. Interestingly, we determined that STIM2 has lower expression in inhibitory than in excitatory neurons (Sup. Fig. 7) – it may therefore be speculated that the roles of STIM proteins are synapse-type specific.

**Figure 6.**
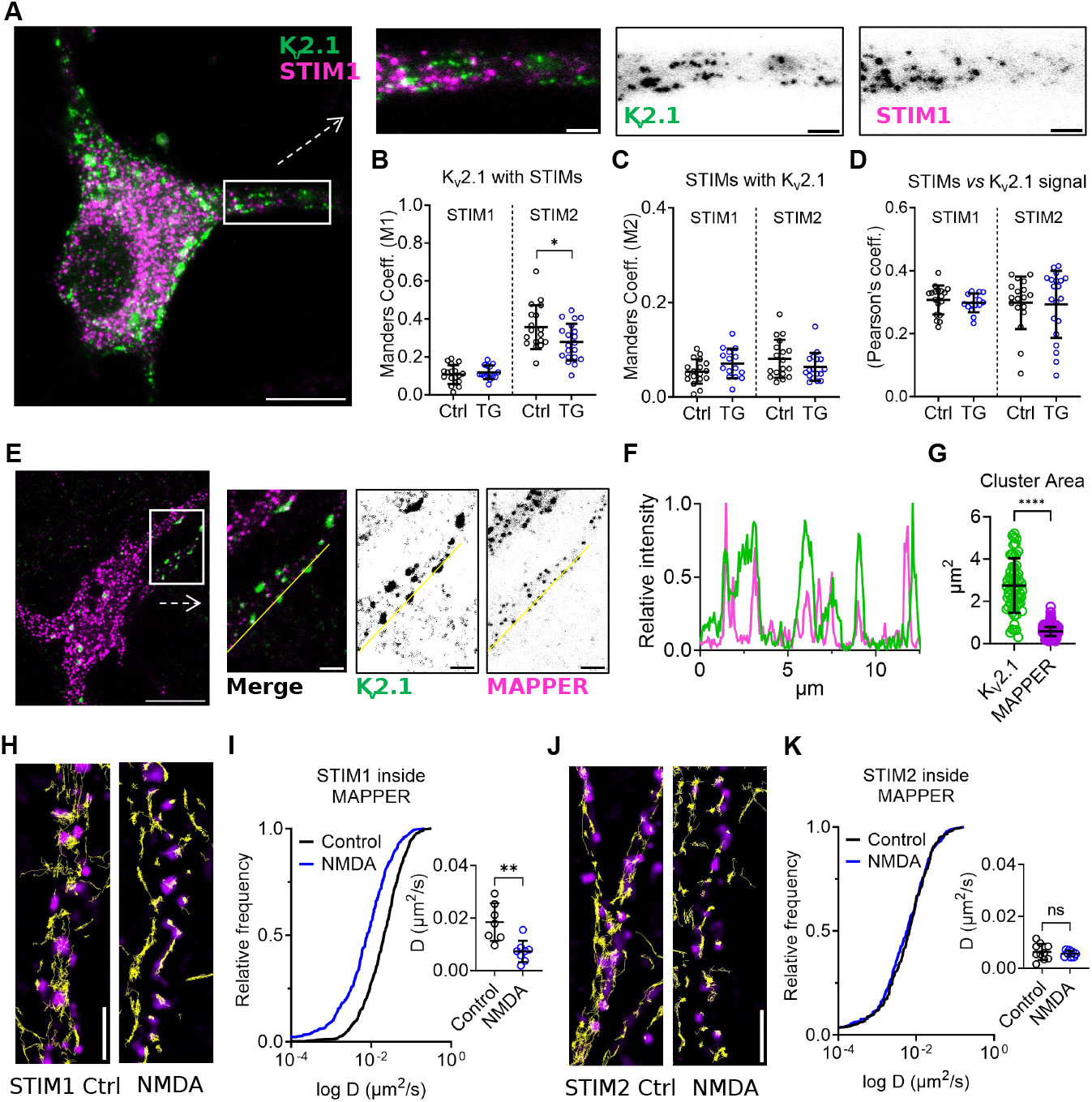
Colocalization and Dynamics of Kv2.1, STIM1, and STIM2 at ER-PM Junctions. **A)** Fixed immunolabeling of Kv2.1 (green) and STIM1 (magenta) under control conditions in hippocampal neurons. **B-D)** Quantification of colocalization between Kv2.1 and STIMs with Manders’ overlap coefficients (*M*_1_ and *M*_2_) and Pearson’s correlation coefficient before and after store depletion induced by 1 µM thapsigargin. **B)** *M*_1_ represents the fraction of the STIMs signal that overlaps with the Kv2.1 signal. **C)** *M*_2_ quantifies the fraction of the Kv2.1 signal that overlaps with the STIMs signal. **D)** Pearson’s correlation coefficient measuring the linear correlation between the STIM1 and Kv2.1 signals. **E)** Fixed immunolabeling of Kv2.1 and MAPPER: representative images of hippocampal neurons immunolabeled for Kv2.1 (green) and MAPPER (magenta). MAPPER-GFP was overexpressed and delivered via rAAV. Scale bars: 20 m for the full image and 2 m for the enlarged section. **F)** Line plot analysis of Kv2.1 and MAPPER signal intensity across the dendritic region. **G)** Size comparison of Kv2.1 and MAPPER clusters from two cultures (10 neurons each). **H-K)** Tracking and Diffusion Analysis of STIM1 and STIM2 at ER-PM Junctions Labeled by MAPPER. Trajectories of STIM1 (**H**) and STIM2 (**J**) molecules (yellow tracks) in control and NMDA-treated conditions, overlaid on MAPPER-labeled ER-PM junctions (magenta). **I, K)** Distribution of diffusion coefficients for STIM molecules within MAPPER spots, comparing control and NMDA-treated conditions. Statistical analysis of STIM1 (**I**) and STIM2 (**K**) diffusion coefficients before and after NMDA treatment, with each data point on scatter plot representing the median diffusion coefficient from a single acquisition encompassing multiple MAPPER spots from one neuron. Statistical significance was determined by parametric unpaired *t*-test and is indicated as ** *P* < 0.01, * *P* < 0.1. Error bars for all figures depict the mean ± SD. Refer to supplementary table for *p*-values, *N, n*, and trajectory counts.

Postsynaptic STIM2 has been proposed to impact on the trafficking of AMPAR subunits to extra-synaptic sites and contribute to plasticity induced increase in synaptic AMPARs (Garcia-Alvarez, Lu, et al., 2015). STIM1 and STIM2 were both shown to colocalize and regulate the activity of AMPA and NMDAR subunits (Gruszczynska-Biegala et al., 2016, 2020; Serwach & Gruszczynska-Biegala, 2020). Our data provide further evidence that NMDA receptors are an important inductor of STIM protein clustering. Stimulation of NMDA caused immobilization and clustering of STIM molecules that was much more robust than ER Ca^2+^ store depletion alone, despite the fact that the magnitude of store depletion was similar in the case of both treatments (**Fig. 4M**). Also, the morphology of STIM clusters that were induced by SERCA blockade and NMDAR stimulation was strikingly different (**Fig. 4J**).

One obvious candidate for such different aggregations would be the interaction of STIMs with lipids of the plasma membrane. The cytosolic, C-terminus of STIMs contains the polybasic domain (PBD), which is responsible for interaction with PM lipids (reviewed in Sallinger et al., 2023). Recent data suggest that the canonical STIM-ORAI activating region (SOAR) is also capable of interacting with lipids (Cohen et al., 2023). Importantly, decreasing the levels of cholesterol in the PM impaired thapsigargin-induced clustering of STIM1 (Pani et al., 2008). Indeed, glutamate-induced clustering of STIM1k (STIM1 with PBD domain removed that impairs its interaction with lipids of the PM) was less strong than that of wild-type STIM1 (**Fig. 4H**). It may be speculated that NMDA activation and passive ER store depletion differentially impact the local organization of lipids in ER PM junctions. Recent data which suggest the interplay between the activity of NMDA, lipids and lipid transporters substantiate this assumption. Sun et al. showed that NMDA stimulation causes dissociation of TMEM24, a lipid transporter, from ER-PM junctions, and that TMEM24 occupied the same population of ER PM contacts as KV2.1 (E. W. Sun et al., 2019). Clusters of both TMEM24 and KV2.1 dissolved upon NMDA stimulation, while in our experiments the same treatment induced a massive clustering of STIMs (**Fig. 5, Sup. Fig. 7**). Ganglioside GM1 is reported to be associated with phosphorylated NMDARs in ER-PM junctions, and GM1 accumulation increased NMDAR currents, leading to spine enlargement (Weesner et al., 2024).

By investigating the molecular players behind the glutamate-induced clustering of STIMs, we confirm that this process is driven by NMDA receptors, while AMPARs or CaV1.2 channels are not involved. This was a surprising result, given the relatively well-established role of STIMs, in particular SITM1, as the inhibitor of CaV1.2 (Dittmer et al., 2017; Park et al., 2010; Pascual-Caro et al., 2018; Y. Sun et al., 2017; Wang et al., 2010). It has to be noted, however, that a study in cardiomyocytes reported an increase in CaV1.2 currents upon STIM1 overexpression, which speaks against this hypothesis (Correll et al., 2015). Data obtained by a different research group showed that overexpression of STIM1 in these cells had no impact on CaV1.2 currents, neither in physiological conditions, nor when the currents were augmented by BayK (Troupes et al., 2017). Similar data were obtained with the use of neuroblastoma cell line, where overexpression of STIM1 had little impact on K^+^ -evoked Ca^2+^ influx into the cytosol (Ramesh et al., 2021). Here, we show that knockdown of STIM1 or STIM2 in dissociated mouse hippocampal neurons did not augment depolarization-induced Ca^2+^ influx, as measured by Fura-2 imaging (**Fig. 5 I, J**). In a heterologous expression system, ER store depletion or STIM1 overexpression failed to attenuate whole cell CaV1.2 currents (**Fig. 5 K, L**). Moreover, inhibition or activation of CaV1.2 by nimodipine and BayK, respectively, did not impact the dynamic behavior of STIMs or the degree of NMDA-induced ER Ca^2+^ depletion (**Fig. 5, Sup. Fig. 4C**). Altogether, these results do not confirm the functional role of STIMs as a direct inhibitor of CaV1.2 channels, or the impact of CaV1.2-dependent Ca^2+^ fluxes on the dynamics of STIMs. This is strengthened by the fact that CaV1.2 was shown to colocalize with KV2.1 in ER-PM junctions (Vierra et al., 2019, 2021), while STIM molecules are only partially colocalized with clusters of KV2.1 (**Fig. 6 A-D, Sup. Fig. 6**). It remains to be determined whether STIMs are implicated in the activity of other voltage-gated Ca^2+^ channels.

Taken together, our results show that neuronal STIM proteins are highly diffusive, and their mobility drastically changes upon various stimuli that modulate neuronal activity. Assessing the distribution of these proteins by means of immunocytochemistry provides only a static snapshot and is susceptible to artifacts. Such artifacts may exaggerate the prevalence of clustered STIM proteins. Whether these clusters are exclusively functional units linked to ORAI channels in the plasma membrane is rather unlikely, since store-operated calcium entry in neurons is very small compared to non-excitable cells (Lu & Fivaz, 2016). NMDARs play a central role in the organization of STIM proteins, but we found no evidence for an interaction between STIM proteins and CaV1.2 channels. While significant differences in the mobility of axonal and dendritic STIM molecules exist, our data suggest that neuronal STIM proteins are primarily active in extra synaptic regions on both pre-and postsynaptic sides. As proposed in a recent publication (Dittmer & Dell’Acqua, 2024), the primary function of STIM proteins may be to regulate the spatial proximity between effector molecules in the plasma membrane and those in the ER membrane. This is confirmed by a very recent study, showing that regularly spaced dendritic contacts between the ER and the PM are organized by junctophilins and STIMs, with implications for long-range dendritic signal integration (Benedetti et al., 2024). Importantly, these contacts were shown to be absent from dendritic spines, confirming our SPT data.

Communication between calcium entry and ER-mediated CICR in glutamatergic spines does not always require perfect colocalization to be functionally effective (Basnayake et al., 2019). However, it is more efficient when the spatial distance between these two membranes is reduced. Therefore, we propose that STIM2 primarily maintains ER-PM contacts, while STIM1 enlarges and strengthens these connections in an activity-dependent manner. This process is reversible even after prolonged NMDAR activation. The different affinities of STIM2 and STIM1 to changes in ER calcium content are well-tuned to dynamically control the extent of ER-PM junctions without the need for additional binding partners in the plasma membrane. This does not preclude the possibility that prolonged confinement of STIM proteins could lead to the trapping of other proteins within ER-PM contacts. These proteins may not directly interact with STIMs but may become confined within the crowded environment of the ER-PM junction, similarly to other specialized membrane areas like pre- or postsynaptic membranes (Choquet & Triller, 2013).

## 4. Author contributions

A.C. and M.H. performed SPT-SMLM microscopy. A.C. optimized and acquired SPT, performed WB, qRT-PCR, vector cloning, and rAAV design and production. F.M. conducted calcium imaging, confocal microscopy and electrophysiological experiments. N.S. created STIM knock-in constructs and optimized the CRISPR-Cas protocol for Knock-ins. A.C., F.M., and M.H. analyzed the data. M.H. provided funding and conceptualized the work, with input from A.C. and F.M. A.C., F.M., and M.H. wrote the manuscript.

We sincerely thank Dr. Anita Heine, Ana Carolina Palmeira do Amaral, Michela Borghi, Corinna Werkmann, and Abderazzaq El Khallouqi for preparing neuronal cultures, as well as Dr. Jennifer Heck, Dr. Junaid Akhtar, and Nathalie Philipp for their lab and experimental support.

## 6. Methods

### Primary hippocampal neuronal cultures

Primary cell cultures were prepared from the following mouse lines: B6.Cg-Stim1^*tm1Rao*^/J (Stim1^*fl/fl*^) and B6.Cg-Stim2^*tm1Rao*^/J (Stim2^*fl/fl*^) OhHora2008; Jackson Laboratory, strain numbers 023350 and 023351, respectively), GAD67-GFP knock-in mice; Tamamaki2003, and Gt(ROSA)26Sor^*tm1*.*1(CAG-cas9*,-EGFP)Fezh*^/J (RRID:IMSR_JAX:024858, Common Name: Rosa26-Cas9 knockin). Dissociated primary hippocampal neurons were obtained from mouse pups (collectively referred to in this study as STIM1, STIM2, Cas9, or GAD67-GFP) housed at the Translational Animal Research Centre (TARC) in Mainz. Hippocampi were dissected from postnatal day 0 (P0) mouse pups in compliance with European and local animal welfare regulations. The tissue was washed three times with cold HBSS (Sigma, H9394) and incubated with 1× trypsin-EDTA (Gibco 15400-054) for 15 minutes to facilitate dissociation. Cells were seeded onto poly-L-lysine (Sigma, P1149) coated Ø18 mm glass coverslips at a density of 70,000 cells per coverslip in 12-well plates. Neurons were cultured in Neurobasal (Gibco, 12348-017) medium supplemented with 2% B27 (Gibco, 17504044) and 5 mM L-glutamine (Gibco, 25030-024) and maintained at 37°C in a humidified 95% air/5% CO_2_ incubator.

### HEK 293T cell culture

HEK 293T cells (Originally referred to as 293tsA1609neo from DSMZ, Ref. ACC 635) were cultured in DMEM (Gibco, 41966-029) supplemented with 10% fetal bovine serum (Thermo, V30160.03) and 1% L-glutamine (Gibco, 25030-024) at 37°C in a humidified 5% CO_2_ atmosphere and passaged weekly. Cells were maintained in T25 flasks and transferred to poly-D-lysine-coated (MERK-Sigma, P1149) Ø18 glass coverslips for experiments. For sub-culturing or experiments, cells were briefly washed with HBSS (MERK-Sigma, H9394) and detached using trypsin-EDTA (Gibco 15400-054), then transferred to 12-well plates at a 1:10 dilution.

### Transfection of cell line

HEK 293T cells in 12-well plates were transfected with 1 g of plasmid DNA using Polyethylenimine PEI (1 mg/ml, Sigma, 408727) at a 1:3 w/v ratio. DNA and PEI were diluted separately in Opti-MEM (Gibco, 31985-070), combined, and incubated for 10 minutes at RT. A 100 l DNA-PEI complex was added to 70% confluent cells, followed by gentle swirling. Media was replaced 3–6 hours post-transfection, and gene expression was analyzed 48–72 hours later.

### Transduction of hippocampal neuronal cultures

For transgenic gene delivery, hippocampal neurons were infected with AAV2 or AAV DJ-6xHis at DIV 2 for experiments involving Cre recombinase-mediated knockdown or at DIV 7 for the introduction of constructs encoding genes of interest (GOIs) such as STIM1, STIM2, ER-GCaMP-150, synaptophysin-GCaMP6f, PSDfingr, or Mapper. The acquisition and analysis of infected neurons were conducted between DIV 7 and DIV 14 to monitor gene expression and the efficiency of transgene delivery. Prior to using each new viral preparation, infectivity assessments were performed using serial dilution to fine-tune the optimal infection volume, ensuring optimal or a 100% infection rate. For specific experiments, lower infection rates were intentionally applied and are detailed within the respective experiment.

### Knockdown of STIMs

To knock-down the expression of STIM1 and STIM2, neuronal cultures set up from Cre/loxP mice (JAX strains no 023350 and 023351, respectively) were transduced with AAV-DJ-hSyn-Cre-P2A-TdTomato virus at DIV2. The efficiency of knockdown was determined by performing immunolabeling at DIV 14-16, followed by confocal microscopy. To achieve moderate transduction efficiency that would enable the comparison of naïve and knock-down neurons on the same coverslip, low concentration of viral particles was used. The intensity of STIM1 and STIM2 signal in TdTomato-positive and negative neurons was compared. As a negative control, neurons were transduced with viral particles coding for a mutated, inactive Cre recombinase (AAV-DJ-hSyn-Cre-P2A-TdTomato).

### Generation of CRISPR-Cas9 STIM1/STIM2 knock-ins

STIM1 and STIM2 knock-ins were generated using the pORANGE template vectors, based on the method described by Willems et al. (2020), with modifications to the knock-in cassettes. These cassettes included the U6 promoter, gRNA, and donor DNA (Halotag in this study) but excluded SpCas9. Instead, floxed Cas9 mouse lines were utilized for Cas9 expression. The modified knock-in cassettes were integrated into an AAV vector and delivered to neuronal cultures at DIV2, along with an AAV vector carrying Cre recombinase for simultaneous transduction. To ensure minimal impact of the inserted tag sequence on protein function, we carefully analyzed the functional domains of STIM1 and STIM2, including signal peptide presence. The tag was positioned close to or immediately following the signal peptide to preserve protein integrity, similar as in our over-expression constructs. Genomic sequences were obtained from the UCSC genome browser to identify suitable PAM sites within these regions. Target sequences were selected with attention to the MIT guide specificity score, ensuring precision in genome editing. The sequences chosen were CCCGCGGAGGTGCCGGGGCA for STIM2 and GTATTCTTTTCACTGTGACT for STIM1.

### Recombinant DNA

Plasmid DNA used in this study included *pMO91_human-STIM1-YFP* (Addgene, ref# 19754; Prakriya et al., 2006), *pEX-YFP-STIM1-deltaK* (Addgene, ref# 18861; Liou et al., 2007), and *pEX-CMV-SP-STIM2*(15-746) (Addgene, ref# 18868; Brandman et al., 2007). Constructs generated or modified within this study included *pAAV_hSyn_SP-Halo-STIM1* (23–685aa) (based on NP_003147.2, NCBI), *pAAV_hSyn_SP-Halo-STIM2* (15–746) (NP_065911.3, NCBI), *pAAV_hSyn_SP-Halo-TMC, pAAV2_hSyn_SP-Halo-STIM1-deltaK* (23–670aa) (NP_003147.2, NCBI), *pAAV-Ef1a-mCherry-IRES-CreMut_Y324F, pAAV-hSyn-MutCre-P2A-dTomato, pAAV_hSyn_synaptophysin-GCaMP6f* (NM_012664.3), *pAAV_CMV_ER-GCAMP-150*, pAAV-DJ-6xHis587, and *pAAV_CMV_GFP-MAPPER*. Additionally, the *pAAV-hSyn-Cre-P2A-dTomato* plasmid was obtained from Addgene (ref# 107738), *pAAV-Ef1a-mCherry-IRES-Cre* (Addgene, ref# 55632), *ER-GCaMP6-150* (Addgene, ref# 86918; de Juan-Sanz et al., 2017), pXX2 (rAAV2-6XHis587), provided by Prof. David V. Schaffer, UC Berkeley (Koerber et al., 2007), *pAAV Helper* (Cell Biolabs), *pAAV2/2* (Addgene, ref# 104963), AAV-GFP/Cre (Addgene, ref# 49056; Kaspar et al., 2002), and *AAV-CMV-GFP* (Addgene, ref# 67634; Xiong et al., 2015).

In this work, the majority of constructs utilized an rAAV-based vector backbone with a human synapsin promoter to drive neuron-specific expression. The synthetic rAAV2 backbone containing ITRs was derived from the *AAV-CMV-GFP* plasmid (Addgene, #67634). For the *pAAV_hSyn_SP-Halo-STIM1* (23–685aa) construct, the SP-STIM1 sequence was amplified from Addgene, ref# 19754, while the Halo tag sequence was amplified from the TUBB-5 Halo plasmid (Addgene, #64691). These elements, along with the hSyn promoter and a (GS4)_2_ flexible linker between the Halo tag and STIM1, were ligated into the rAAV backbone. Similarly, for the *pAAV_hSyn_SP-Halo-STIM2* construct, the STIM2 sequence was amplified from Addgene, ref# 18868. For the *pAAV_hSyn_SP-Halo-TMC* construct, *pAAV_hSyn_SP-Halo-STIM1* was used as the template, with the STIM1 sequence removed and the transmembrane (TM) region retained immediately after the Halo tag sequence. This construct, which included the SP (1–23aa) of STIM1 followed by the Halo tag and the STIM1 transmembrane region, served as a Halo tag control. The *AAV2_hSyn_SP-Halo-STIM1-deltaK* construct was created by amplifying the STIM1-deltaK fragment from Addgene, ref# 18861, and replacing the STIM1 sequence in *pAAV_hSyn_SP-Halo-STIM1* with the STIM1-deltaK fragment. For Cre-recombinase control experiments, the *pAAV-Ef1a-mCherry-IRES-CreMut_Y324F* and *pAAV-hSyn-MutCre-P2A-dTomato* constructs were generated by inserting the Y324F mutation from pAAV-Ef1a-mCherry-IRES-Cre (Addgene, ref# 55632) into these plasmids, using pAAV-hSyn-Cre-*P2A-dTomato* (Addgene, ref# 107738) as a reference. The *pAAV_CMV_ER-GCAMP-150* construct was created by amplifying the *CMV_ER-GCAMP-150* sequence from Addgene, ref# 86918, and inserting it into the rAAV backbone. Lastly, *pAAV-DJ-6xHis587* was designed following the method described by Koerber et al., 2007, where a 6-residue histidine tag was inserted at the 587 aa position in the *pAAV2-N587X* plasmid (Addgene, #130877).

### Antibodies and Chemicals

Primary antibodies and dyes used in this study were purchased from the following sources: Bassoon (Synaptic Systems, 141 004, 141021), Beta-actin (Synaptic Systems, 251 011), Homer 1 (Synaptic Systems, 160011, 160003), RIM (Synaptic Systems, 140203), Ankyrin-G (Neuromab, N106/36), Kv2.1 (NeuroMab, 75-159), MAP2 (Synaptic Systems, 188011, 188004), NR2B (Alomone Labs, AGC-003), STIM1 (Sigma-Aldrich, S6072), STIM2 (Alomone Labs, ACC-064), and Halo-Tag 646 (Promega, G9281), Fura-2 AM (abcam, ab120873), HRP anti-Guinea Pig (Jackson IR, 706-035-148) and HRP anti-Mouse (Jackson IR, 115-035-146).

Pharmacological agents used in this study: DHPG (HelloBio, HB0045), Bicuculline Methiodide (Tocris, 2503), Caffeine (Sigma-Aldrich, C0750), CNQX Disodium Salt (HelloBio/Tocris, HB0204/1045), Cyclopiazonic Acid (CPA) (HelloBio, HB1117), D-AP5/D-APV (HelloBio/Tocris, HB0225/0106), Ifenprodil (HelloBio, HB0339), L-Glutamate (abcam, ab120049), MK 801 Maleate (HelloBio, HB0004), Nimodipine (Tocris, 0600/100), NMDA (N-Methyl-D-Aspartic Acid) (HelloBio, HB0454), Ryanodine (HelloBio, HB1320), and TG (Thapsigargin) (HelloBio, HB1118).

### HaloTag Labeling

HaloTag labeling was performed using cell-permeable Halo-JF646 (Promega, G9281) ligand. The ligands were prepared as 200 *μ*M stock solutions in water and stored at 20°C in single-use aliquots. For live-cell single-particle tracking (SPT), HaloTag ligands were added to the culture medium at a final concentration of 1 nM, and cells were incubated for 30 minutes to allow labeling before SPT imaging and data acquisition. For fixed staining, 20 nM of the ligand was incubated with cells for 3 hours. After incubation, cells were washed twice with neurobasal medium for 5 minutes each, followed by fixation and permeabilization.

### AAV Production

A helper virus-free system was used for recombinant adeno-associated virus (rAAV) production. HEK293T cells were cultured to 70% confluency in a T75 flask and transfected with a 1:2 (w/v) plasmid DNA to PEI (1mg/ml) ratio. For transfection, Equimolar amounts of three plasmids (helper, viral coat—either AAV2 or DJ6xHIS—and the gene of interest) were combined for a total of 30 *μ*g per flask and incubated in 0.9 ml Opti-MEM (Gibco, 31985-070) for 10 minutes. The transfection mixture was then added to the cells and incubated for 6 hours at 37°C with 95% air and 5% CO2, followed by media replacement with fresh DMEM (Gibco, 41966-029) supplemented with 10% fetal bovine serum (Thermo, V30160.03) and 1% L-glutamine (Gibco, 25030-024). After 48 hours, cells were scraped and harvested in 1 ml ice-cold resuspension buffer (50mM Tris; 1M NaCl; 0.001% Pluronic (10%)), subjected to three freeze-thaw cycles, and centrifuged at 13,000g for 15 minutes at 4°C to separate the crude viral lysate from cell debris. The crude lysate was purified using an iodixanol density gradient and ultracentrifugation. Sequential layers of 15%, 25%, 40%, and 54% iodixanol were added to quick-seal tubes (Beckman Coulter, 342414) containing 7 ml of viral supernatant, then centrifuged at 350,000g for 2 hours at 18°C under vacuum. The 40% iodixanol layer containing the virus was carefully extracted, and buffer exchange (1mM MgCl2; 2.5mM KCl; 0.001% Pluronic (10%); pH 7 in 1xPBS) was performed using Amicon ultra-15 filters (100 kDa Millipore, UFC910024), concentrating the volume to 250 *μ*l at 3000 RPM at 4°C. The purified rAAV was stored at -80°C in single-use aliquots.

### Western Blotting

Cultured floxed STIM1/STIM2 hippocampal neurons were seeded in 12-well plates and infected at DIV2 or DIV7 with rAAV2 or rAAV DJ-6xHis expressing Cre recombinase to achieve 100% infection (verified through in-lab titration). Cells were harvested at DIV14–16 and lysed in 100 *μ*l SDS sample buffer (62.5 mM Tris-HCl (pH 6.8), 2% SDS, 10% Glycerol, 0.001% Bromophenol Blue, 5% *β*-Mercaptoethanol) per well, with samples pooled when needed. Protein concentration was measured using the Bradford assay, and 20 *μ*g of protein was denatured at 95^°^C for 5 minutes before SDS-PAGE. Proteins were separated on a 10% resolving gel (10% Acrylamide (Rotiphorese Gel 40, Roth, 3030.3), 0.375 M Tris-HCl (pH 8.8), 0.1% SDS, 0.1% Ammonium Persulfate, TEMED) following a 5% stacking gel (5% Acrylamide, 0.125 M Tris-HCl (pH 6.8), 0.1% SDS, 0.1% Ammonium Persulfate, TEMED) at 8 mA per gel for stacking and 12 mA per gel for resolving. The PVDF membrane (0.45 *μ*M, Roth, T830.1), activated with methanol (Roth, 7342.1), was used for transfer in cold buffer (20% methanol) at 4^°^C for 3 hours at 200 mA. Membranes were washed with TBST (20 mM Tris-HCl, 150 mM NaCl, 0.1% Tween-20) and blocked with 5% non-fat milk in TBST for 1 hour at room temperature.

Primary antibodies (1:1000 in 5% milk TBST, STIM1 (N-Terminal), Sigma-Aldrich, S6072; and STIM2, Alomone labs, ACC-064) were applied overnight at 4^°^C, followed by three TBST washes. Secondary HRP-conjugated antibodies (all from Jackson IR: HRP anti-Guinea Pig, 706-035-148; HRP anti-Mouse, 115-035-146; HRP anti-Rabbit, 711-035-152) (1:2000–1:5000 in 5% milk TBST) were incubated for 1 hour, then membranes were washed twice for 10 minutes each. Protein bands were visualized with HRP substrate (Pierce ECL Western Blotting Substrate, Life Technologies, 32209) following the manufacturer’s protocol using the Intas ChemoStar imaging system (5-minute acquisition), and band intensities were quantified with ImageJ, normalized to *β*-actin (Synaptic Systems, 251011) as a loading control.

### qRT PCR

Brain tissues (hippocampus, cerebellum, and cortex) were collected from three adult female (3-month-old) STIM1 floxed mice, and up to 100 mg of each tissue was homogenized in 1 mL Trizol (Thermo Fisher, 10296010). After a 5-minute incubation at room temperature (RT), 200 *μ*l chloroform:isoamyl alcohol (24:1) (Sigma, C0549) was added, followed by vortexing for 30–60 seconds. The mixture was incubated at RT for 10–12 minutes and centrifuged at 12,000 g for 15 minutes at 4^°^C, and the upper phase containing RNA was collected and mixed with 700 *μ*l isopropanol (Sigma, 190764). After a 10-minute incubation at RT (or overnight at −20^°^C or −80^°^C for higher RNA yields), RNA was pelleted by centrifugation at 12,000 g for 20 minutes at 4^°^C, washed with 750 *μ*l 75% ethanol (Sigma, 493546), and air-dried for 15 minutes. The pellet was resuspended in 30–50 *μ*l RNase-free water (Qiagen, 129114), incubated at 65^°^C for 5 minutes, and stored at −80^°^C.

RNA (30 *μ*l) was treated with RNase inhibitor (NEB, M0314L) and DNAse I, RNase-free (1 U/*μ*L) (Thermo Fisher, EN0521) at 37^°^C for 20–30 minutes, and purified using the RNeasy Mini spin column (Qiagen, 79523). Reverse transcription (with SuperScript^TM^ III, Invitrogen, 18080051) of 2 *μ*g of RNA was performed in a 20 *μ*l reaction using random/oligo dT primers, incubated at 65^°^C for 5 minutes, followed by cooling on ice for 2 minutes. The remaining RT components were added, and the reaction was incubated at 42^°^C for 1 hour, then heat-inactivated at 80^°^C for 5 minutes, following the manufacturer’s protocol.

For qPCR, 10 *μ*l reactions were set up with SYBR Green dye (Life Technologies, 4367659) using splice variant-specific primers for STIM1 and STIM2, with each reaction performed in triplicate using the ViiA^TM^ 7 Real-Time PCR System (Applied Biosystems). Relative quantification (RQ) was determined using the comparative CT method, with RQ calculated as 2^−Δ*CT*^, where Δ*CT* is the difference between the target gene and housekeeping gene CT values.

### Cytosolic Ca^2+^ imaging

Single-cell somatodendritic cytosolic Ca^2+^ levels were quantified using the ratiometric Ca^2+^ indicator, Fura2 acetoxymethyl ester (Fura-2 AM, abcam, ab120873). Cultured neurons aged 14-16 DIV were loaded with 2 *μ*M Fura2-AM for 20 min at 37°C in Ringer’s solution (NaCl 130 mM, KCl 3 mM, MgCl_2_ 1 mM, CaCl_2_ 2 mM, Glucose 10 mM, HEPES 10 mM, pH 7.4, Osmolarity ∼310 mOsm). After washing with Ringer’s solution, the cells were left undisturbed for an additional 10 min at 37°C to ensure dye de-esterification and equilibration. Acquisition was performed under the IX71 Olympus microscope equipped with a 40x/1.4NA objective. Fura-2 was excited at 340 and 380 nm with a Polychrome 3000 lamp (Agilent Technologies), and emitted light was captured by an EM-CCD camera iXON Ultra 897 (Andor, pixel size: 16 *μ*m × 16 *μ*m, chip size 8.2×8.2 mm). Ca^2+^ levels were calculated as the ratio of fluorescence emitted upon 340 nm excitation to fluorescence emitted upon 380 nm excitation (peak emission at 510 nm at both excitation wavelengths, excitation filters used: 340/26 and 387/11). Background fluorescence was subtracted, signal calibration was not performed. The intracellular Ca^2+^ levels in individual neurons were measured using MetaMorph imaging software (version 7.8.8.0). Microsoft Excel and GraphPad Prism 9 software were utilized for data processing. For recording basal cytosolic Ca^2+^ levels, neurons were placed in Ringer’s solution with 2 mM CaCl_2_ for 2 min. To evaluate VGCC activity and exclude glial cells from analysis, the PM was depolarized by the application of Ringer’s solution containing 50 mM KCl and 10 mM CaCl_2_ (final concentration). The baseline was calculated for the first 20 s, and the magnitude of depolarization-induced Ca^2+^ influx was measured as the amplitude following KCl addition.

### ER Ca^2+^ imaging

Single-cell somatodendritic ER Ca^2+^ levels were monitored using the genetically encoded probe ER-GCaMP6-210 (Addgene no. 86919), which was delivered via an AAV2-based vector (AAV-hsyn-ER-GCaMP150) seven days before the experiment. Cultured neurons aged 14-16 DIV were imaged at 37°C in Ringer’s solution. Acquisition was performed on the IX71 Olympus microscope equipped with a 40x/1.4NA objective. The fluorescent dye was excited with a Polychrome 3000 lamp (Agilent Technologies) and captured by an EMCCD camera iXON Ultra 897 (Andor, pixel size: 16 *μ*m × 16 *μ*m, chip size 8.2×8.2 mm) using MetaMorph imaging software (version 7.8.8.0). 470/495 excitation and 525/50 emission filters were used. Data processing was conducted using Microsoft Excel and GraphPad Prism 9 software. To assess the magnitude and kinetics of depletion of Ca^2+^ from the ER upon passive leak and NMDAR activation, CPA (HelloBio, HB1117) or NMDA (HelloBio, HB0454) were added at t = 100 s. The magnitude of ER Ca^2+^ efflux was determined as the negative amplitude at t = 400 s, and the kinetics were evaluated by fitting the curve with a single exponential function (between t = 100 s and t = 400 s).

### HEK 293 Cav1.2 cell culture for electrophysiology

HEK293T cell line stably expressing full-length human Cav1.2 e^1*b*/8*b*/−9*/22/32/33^ and auxiliary subunits human Cav*α*2*δ*1 and human Cav*β*3 (Oertner et al. 2017) were cultured in DMEM (Gibco, 41966-029) supplemented with 10% fetal bovine serum (Thermo, V30160.03) and 1% L-glutamine (Gibco, 25030-024) at 37°C, and maintained in selective medium (hygromycin 50 *μ*g/ml, blasticidin 15 *μ*g/ml, geneticin 500 *μ*g/mL) with a humidified atmosphere containing 5% CO_2_ and 95% air. For cell detachment during experiments or sub-culturing, the cells were initially washed with HBSS (Sigma, H9394) and subsequently detached using Trypsin + 1x EDTA (Gibco, 25300-062) under sterile conditions. Culturing was carried out in T25 flasks, and for electrophysiological experiments, cells were transferred to poly-D-lysine-coated (MERK-Sigma, P1149) Ø18 glass coverslips (12-24 h before experiments). For STIM1 overexpression experiments, the cells were transfected with a plasmid coding for human STIM1 fused with YFP (Addgene, plasmid no. 19754) 3 days before experiments. Expression of Cav1.2 subunits was triggered by 1 *μ*g/ml doxycycline treatment (applied to the cell medium 2 days prior to experiments).

### Patch-clamp

Pipettes were pulled with a vertical puller PC-100 (Narishige) from borosilicate glass (Scientific Products, 0.86 mm inner diameter, 1.50 mm outer diameter). Resistance of the pipette tip was between 2 and 5 MΩ when filled with intracellular solution that contained 140 mM CsCl, 3 mM MgCl_2_, 0.66 mM CaCl_2_, 10 mM HEPES, 11.7 mM EGTA, 2 mM Na_2_-ATP, 0.3 mM Na_3_GTP. Osmolarity was adjusted with sucrose to 300 ± 5 mOsm/kg H_2_O and pH was set with CsOH to 7.25 ± 5 at room temperature. Extracellular solution contained 140 mM NaCl, 10 mM BaCl_2_, 10 mM glucose, 10 mM HEPES, and 1 mM MgCl_2_. Osmolarity was adjusted with sucrose to 300 ± 5 mOsm/kg H_2_O and pH was set with NaOH to 7.40 ± 5 at room temperature. In ER Ca^2+^ store-depletion experiments, thapsigargin was added to the Ringer solution 10 min before the start of experiments (final concentration 1 *μ*M). In control conditions, DMSO (1:1000 v/v) was used. Recordings were performed at room temperature. After establishment of the whole-cell mode, the cells were voltage-clamped at -60 mV and access resistance was compensated so that the value did not exceed 5 MΩ. To activate the Cav1.2 channels, voltage steps from -50 mV to 60 mV were applied every 5 mV. Leak currents have been subtracted online before the channel activation protocol. Signals were acquired with Patchmaster software (HEKA Elektronik), filtered on-line with a 3 kHz low-pass filter, sampled at 20 kHz, amplified by the EPC9 amplifier and digitized with the LIH 8+8 AD/DA converter (HEKA Elektronik). Inward current amplitude was quantified (Fitmaster, HEKA), plotted, and analyzed for statistical significance (Prism, GraphPad).

### Immunolabeling and Confocal Microscopy

Hippocampal cultures from DIV 14-16 were used for immunolabeling. In a subset of experiments, passive ER Ca^2^+ store-depletion or glutamate stimulation were performed immediately before fixation. The cells were fixed by adding 500 µl of 4% paraformaldehyde (PFA) onto each Ø18 coverslip and incubating for 10 min at 37°C. Next, the 4% PFA solution was removed, and permeabilization was performed by adding 0.5 ml of permeabilization buffer (0.3% Triton in 1xPBS (83 mM Na_2_HPO_4_, 17 mM NaH_2_PO_4_, 150 mM NaCl)) for 10 min at RT. Following permeabilization, the coverslips were washed three times for 10 min with washing/blocking buffer that was composed of 2% albumin fraction V (ROTH, CAS no. 9048-46-8), 25 mM glycine, 1x caseine (Sigma-Aldrich, cat. no. B6429) at RT. For immunostaining, coverslips were incubated with primary antibodies targeting the protein of interest diluted in washing/blocking buffer (1:200 dilution) for 4 hours at RT or overnight at 4°C, and then washed three times with washing/blocking buffer. Subsequently, coverslips were incubated with secondary antibodies diluted in washing/blocking buffer (1:1000 dilution) for 1 h at RT in the dark and washed again three times. The final washes were performed twice with 1x phosphate-buffered saline (PBS), each for 5 min at RT. Finally, coverslips were mounted with 10µl of Mowiol® mounting medium (Roth, 713.2). The mounting medium was allowed to harden overnight at RT in the dark before further analysis. Acquisition of images was performed under Leica Stellaris8 confocal microscope, using 100x 1.4 NA oil objective, white light laser and HyD detectors, controlled by LasX software. Pixel size was 60 nm, Z-stacks were acquired with an axial resolution of 200 nm. Images were processed with ImageJ software (NIH).

### SPT Acquisition and Analysis

Imaging experiments for SPT of Halo tagged STIMs were performed on a Nikon Eclipse Ti 2 E inverted microscope equipped with an oil immersion TIRF objective (60 x, NA 1.45 expanded by 1.5 x magnification lens) and the perfect focus system for Z focus stabilization. Samples were illuminated in TIRF mode, and images were obtained with an exposure time of 50 ms for 500 frames, 20 Hz using an Orca fusion BT sCMOS camera (Orca Flash 4 0 Hamamatsu Photonics, pixel size: 6.5 x 6.5 µm, chip size 13.3 x 13.3 mm). The imaging acquisition was controlled by the NIS Elements advanced research acquisition software (Nikon). During the acquisition, the fluorophores (GFP and Halo tag jenelia 646) were excited by 488 nm and 640 nm at 20% laser power (Coherent; MPB Communications Inc) and a 1.5 x lens in the TIRF coupling was used to illuminate the center of the field of view and to focus the laser power, resulting in a pixel size of 0.072 x 0.072 µm^2^. The imaging experiments were carried out at 37°C and perfused with Ringer’s solution for recordings longer than 5 minutes to maintain physiological conditions. Acquired image stacks were analyzed using the PALMTracer software package (Butler et al., 2022) for MetaMorph (version 7.10.3.297). Collaboration with Dr. J.B. Siberita at IINS Bordeaux (France) facilitated the analysis. Spots were detected by thresholding the images and localized by fitting a 2D Gaussian function with a 9-pixel radius that matches the microscope’s point spread function and an initial sigma of 1.6, achieving 40 nm localization accuracy in both x and y dimensions. The average diffusion coefficient (D) for each tracked particle was estimated from the initial MSD versus t curve slope (D = MSD/4t) using the first 8 time points (50–400 ms).

### Workflow of SPT experiment

At DIV 7, Halo-tagged STIMs and synaptic markers were introduced via rAAV infection, with imaging conducted at DIV 14–16. Prior to acquisition, cells were incubated with 646-halo-ligand for 3 hours, followed by pharmacological treatment (for test conditions) and washing steps. Imaging was performed in Ringer’s solution at 37^°^C. For single-particle tracking (SPT) of Halo-tagged STIMs, 500 frames were captured at 20 Hz, while 50 frames were acquired sequentially for the synaptic marker under the same TIRF conditions. Image stacks for both the synaptic label and Halo-tagged STIMs were processed in MetaMorph for ROI selection for localizations. Synaptic ROIs (∼160 nm radius, 12 pixels) were marked by briefly aligning the average of synaptic label frames with the maximum projection of 500 SPT frames. Similar size ROIs were placed in the extra-synaptic region for area normalization. Using the same aligned image, a cell mask was drawn manually to filter out localizations from axons and dendrites (mapper spots were detected by thresholding the images).

Subpixel localizations and trajectory calculations for each region were performed using PALMTracer. To ensure accurate trajectory connections, a maximum fitting radius of 4 pixels (52 nm) between frames was set, based on our visual inspection of the fastest moving molecules in our acquisitions. Localizations exceeding this range were excluded as false connections. The diffusion coefficient was calculated from the mean squared displacement (MSD) and served as a primary metric for data interpretation across experiments. Diffusion coefficients derived from tracks lasting at least 8 frames (400 ms) were included in the analysis, while shorter tracks were excluded. GraphPad Prism9 was utilized for data visualization, with two types of graphs presented in this study. First, diffusion coefficients were shown as cumulative frequency distributions on a logarithmic scale to pool data from identical conditions. Second, to ensure consistent weighting across experiments, a scatter plot of the median diffusion coefficient from each acquisition was plotted and used in subsequent statistical analysis.

