## Supplementary figures and tables for "Activity-Dependent Localization and Heterogeneous Dynamics of STIM1 and STIM2 at ER-PM contacts in Hippocampal Neurons"

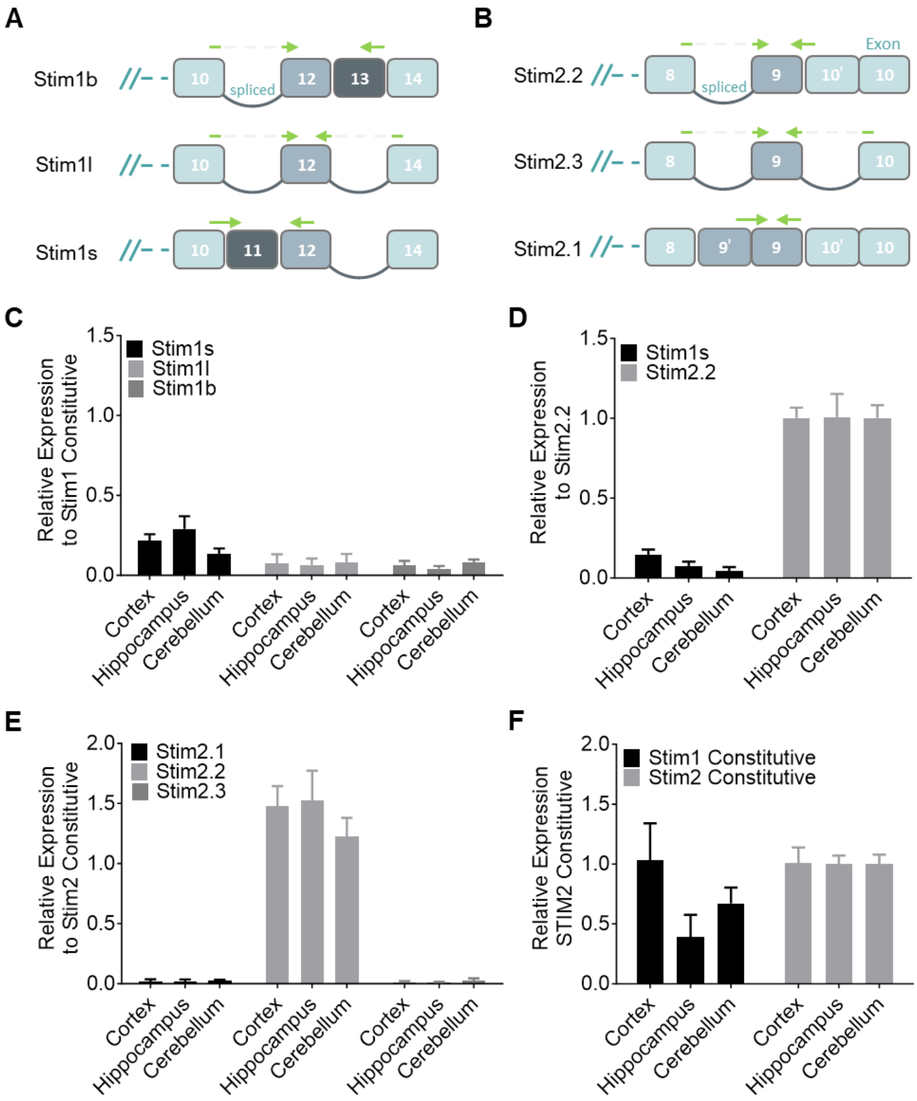

**Sup. Fig.1:** Identification of most abundant Stim1 and Stim2 splice variants in different brain regions. **A)** Schematic representation of Stim1 splice variants, displaying tail axons 10, 11, 12, 13, and 14, along with the locations of the qRT-PCR primer pairs designed for their detection. **B)** Schematic representation of Stim2 splice variants' tail axons 8, 9 and 10, and the corresponding locations of the designed qRT-PCR primer pairs (schematic on Fig. Sup. 10) for their detection. **C)** Normalized expression levels of Stim1 splice variants with respect to the constitutive expression of Stim1. **D)** Comparison of highly expressed Stim1 splice variants with a highly expressed Stim2 splice variant. Relative expression of Stim1s is normalized to Stim2.2. **E)** Relative expression of Stim2 splice variants, emphasizing the predominant expression of Stim2.2 when normalized to the constitutive expression of Stim2. **F)** Comparative analysis of global constitutive expression between Stim1 and Stim2. Stim1 levels are normalized to STIM2 in the respective brain regions. All the experiments were performed in three independent biological replicates, error bars indicate the mean ± SEM. Refer to supplementary table for p-values.

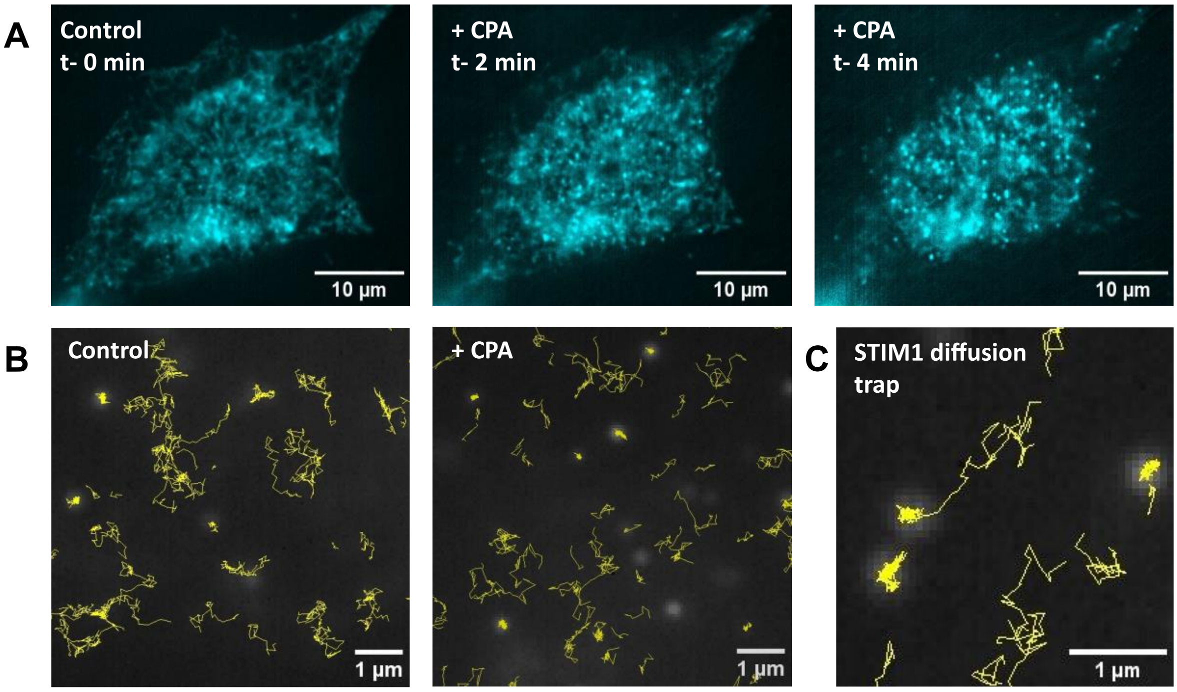

**Sup. Fig.2:** Single particle tracking analysis of Halo-STIM1 in HEK293T cells. **A)** Live cell imaging of HEK293T cells overexpressing Halo-STIM1, labeled with janelia646 halo ligand, and subjected to 10 µM CPA treatment to induce store depletion. Notably, after a 2-minute incubation with CPA, distinctive STIM puncta emerged at the junction between the ER-PM junction. **B)** SPT trajectories of Halo-STIM1 in HEK293T cells in both resting and store depleted conditions. **C)** A visual representation of the diffusion trap model of STIM1 during activation visualized by STIM1 trajectories. Initially mobile, STIM1 molecules are trapped and clustered in ER-PM junction.

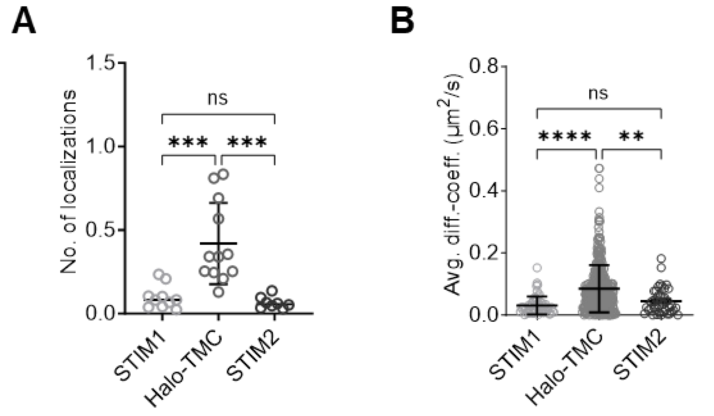

**Sup. Fig.3: A)** Quantification of STIMs' occurrence in the post-synaptic compartment compared to the Halo-TMC control, determined from ROIs surrounding the PSD label. **B)** Corresponding diffusion coefficients of STIMs calculated from ROIs encircling PSD labels.

To obtain a quantitative measure of the occurrence of STIMs in reference to the control, the localizations of STIMs within ROIs encircling the PSD95 label were calculated. ROIs of variable diameters were constructed by the MetaMorph program using an intensity-based approach which included all PSD95 labels, both those located laterally to the dendritic shaft and those on the shaft, without differentiation. Data was collected from over 3000 PSD labels for STIM1, STIM2, and Halo-Control. This data was then normalized based on the number of detected ROIs in each acquisition and plotted as the number of localizations per PSD. This can be interpreted as the frequency of STIM occurrences for each PSD. To improve localization detection, PALMTracer program settings were adjusted to 2 frames (100 ms). This adjustment allowed for accounting for nearly every particle that interacted with the label without missing a significant fraction. With the same settings applied to all constructs, the number of detected Halo-TMC localizations were significantly higher compared to both STIM1 and STIM2, as expected (Sup.Fig.3A). However, it is crucial to interpret this observation cautiously as STIM1 and STIM2 data primarily includes false-positive colocalization events from PSD labels on the dendritic shaft. In contrast, for Halo-TMC, the localizations are derived equally from PSDs located on the lateral and shaft regions. Furthermore, examining the diffusion coefficients calculated from the 100 ms time frame in Sup.Fig.3B clearly demonstrates that the Halo-control exhibits high diffusivity. Consequently, using smaller ROIs for detecting the Halo-control (which exhibits faster movement) would lead to the exclusion of a substantial portion of the data, as the rapidly moving particles would exit the ROI before the 100 ms detection window is completed. Thus, the number of localizations in Sup.Fig.3A is expected to be even higher for Halo-TMC. Conversely, for STIM1 and STIM2, the detected molecules predominantly arise from the PSD marker on the dendritic shaft. As a result, the actual fraction representing the presence of STIMs in post-synaptic compartments is exceedingly low. From the quantification, an average of 0.5 localizations of Halo-TMC were observed for each PSD examined. In contrast, the data for STIM1 and STIM2 indicate a significantly lower range of 0.05 to 0.1 localizations for each PSD, primarily originating from the dendritic shaft. In summary, it can be concluded that STIMs were not present in spines and the post-synaptic compartment, and no enrichment of STIMs in the post-synaptic region was observed. Refer to supplementary table for p-values, N, n, and trajectory counts.

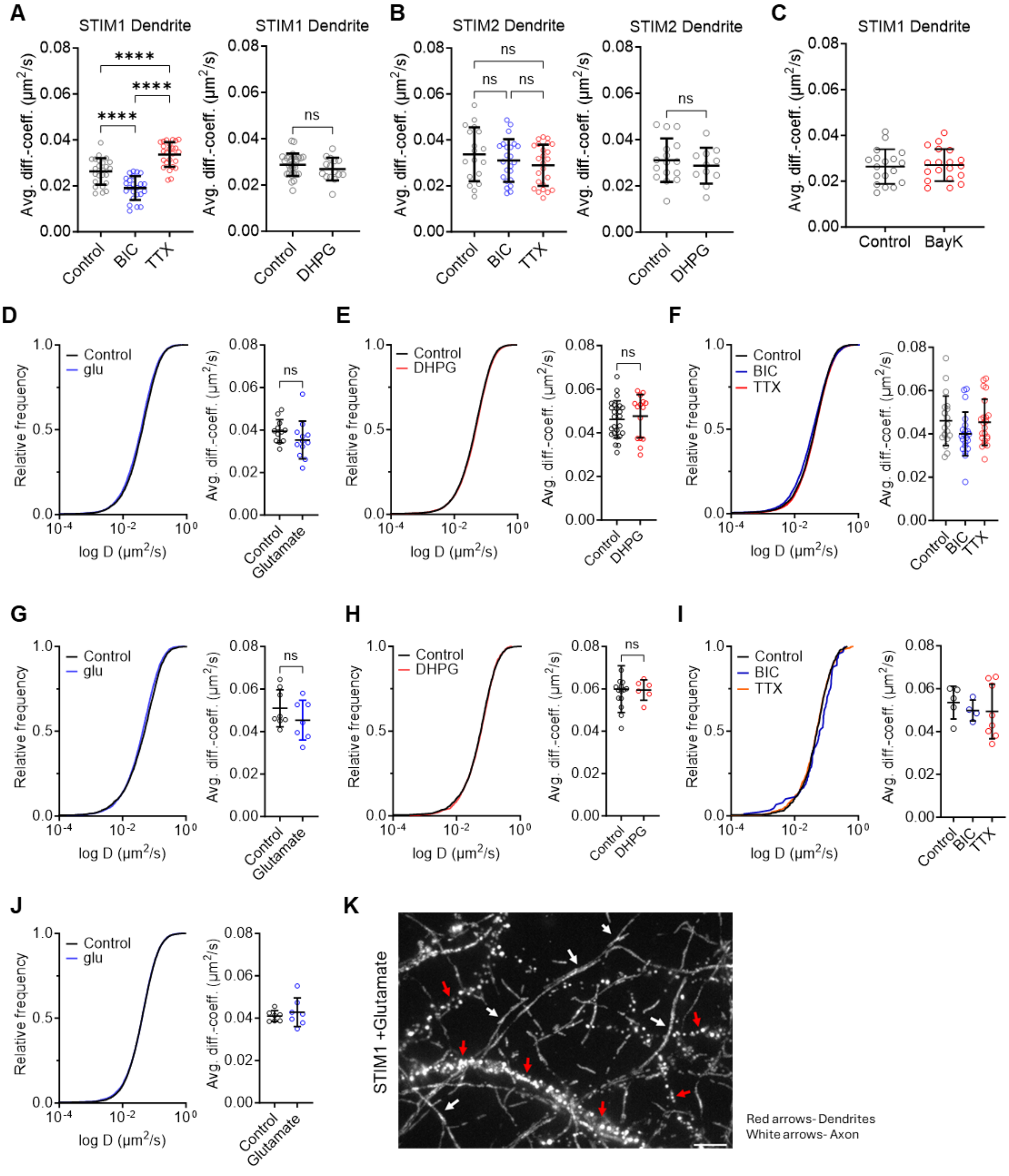

**Sup. Fig.4:** Effect of homeostatic treatments on STIM1 (**A**) and STIM2 (**B**) dynamics in dendrites after 24h treatment with either BIC 10 μM or TTX 2 μM. **C**) Effect of LTCCs action on STIM1 dynamics in dendrite. Cumulative frequency distribution and median diffusion coefficient comparison of STIM1 within dendrites under control (unstimulated) and Bay K-treated conditions (10 μM, LTCC agonist). Each data point on the dot plot represents the median diffusion coefficient from a single acquisition (n = 18 cells for control, n = 19 cells for Bay K treatment, from two independent cultures, N = 2). Experiments were conducted in Ringer's solution at 37°C, containing 2 mM $Ca^{2+}$, 2 mM $Mg^{2+}$, 10 μM CNQX, and 10 μM APV, with or without the addition of 10 μM Bay K. **D, E, F)** STIM1 dynamics in axons in various treatments. **G, H, I)** STIM2 dynamics in axons in various treatments. J) STIM1ΔK dynamics in glutamate treatment. **K)** Max projection of STIM1 SPT movie after glutamate treatment highlighting differences in the cluster appearance of STIM1 in axons and dendrites marked by arrows. All experiments were conducted in a Ringer's solution at 37 °C with the presence of 2 mM $Ca^{2+}$ 2 mM $Mg^{2+}$, 10 μM CNQX, and 10 μM APV. Error bars indicate the mean ± SD. Refer to supplementary table for p-values, N, n, and trajectory counts.

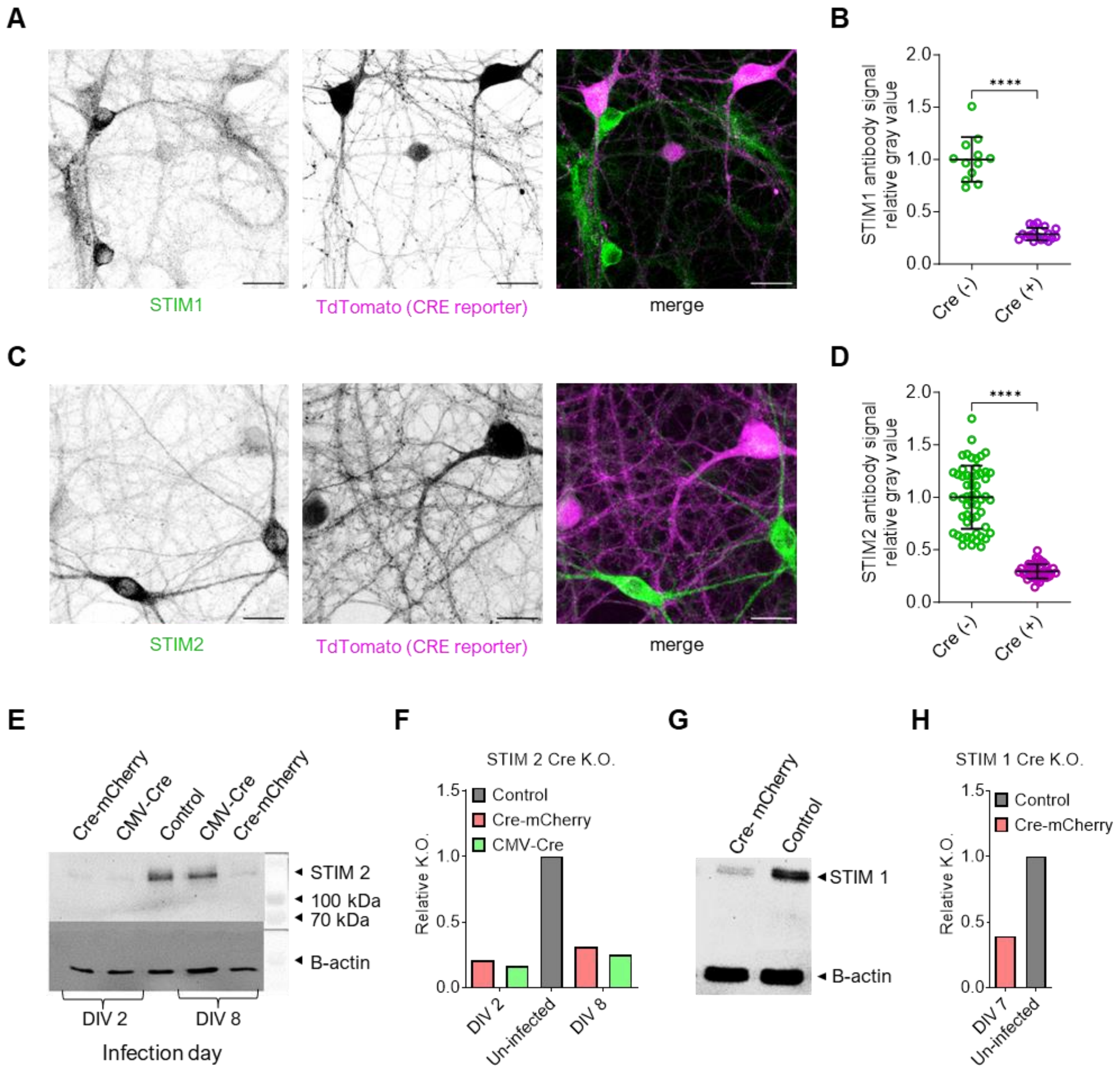

**Sup.**

**Fig.5:** Cre induced knockout of Stims from loxP mouse lines. **A)** Antibody staining of endogenous Stim1 in cultures infected with low level Cre-tdTomato on DIV-1-3 and acquired on DIV 14-16, scale bar: 20µm. **B)** Expression levels (calculated from intensity) of Stim1 in Cre positive and Cre negative neurons. **C)** Antibody staining of endogenous Stim2 in cultures infected with low level Cre-tdTomato on DIV-1-3 and acquired on DIV 14-16, scale bar: 20µm. **D)** Expression levels (calculated from intensity) of Stim2 in Cre positive and Cre negative neurons. **E)** Western blot from floxed Stim2 hippocampal neurons infected with rAAV Cre recombinase (Cre-mCherry and CMV_Cre-GFP) at DIV2 and DIV7. Cells harvested on DIV14, 20 µg of total protein detected with rb-anti-Stim2 Alomone antibody and anti-β-actin, developed on same bolt. **F)** Signal quantification from blot. **G &** **H**) Western blot from floxed Stim1 hippocampal neurons infected with rAAV Cre-mCherry at DIV7. Cells harvested on DIV14, 20 µg of total protein detected with rb-anti-Stim1 Sigma antibody and anti-β-actin, developed on same bolt. Refer to supplementary table for p-values.

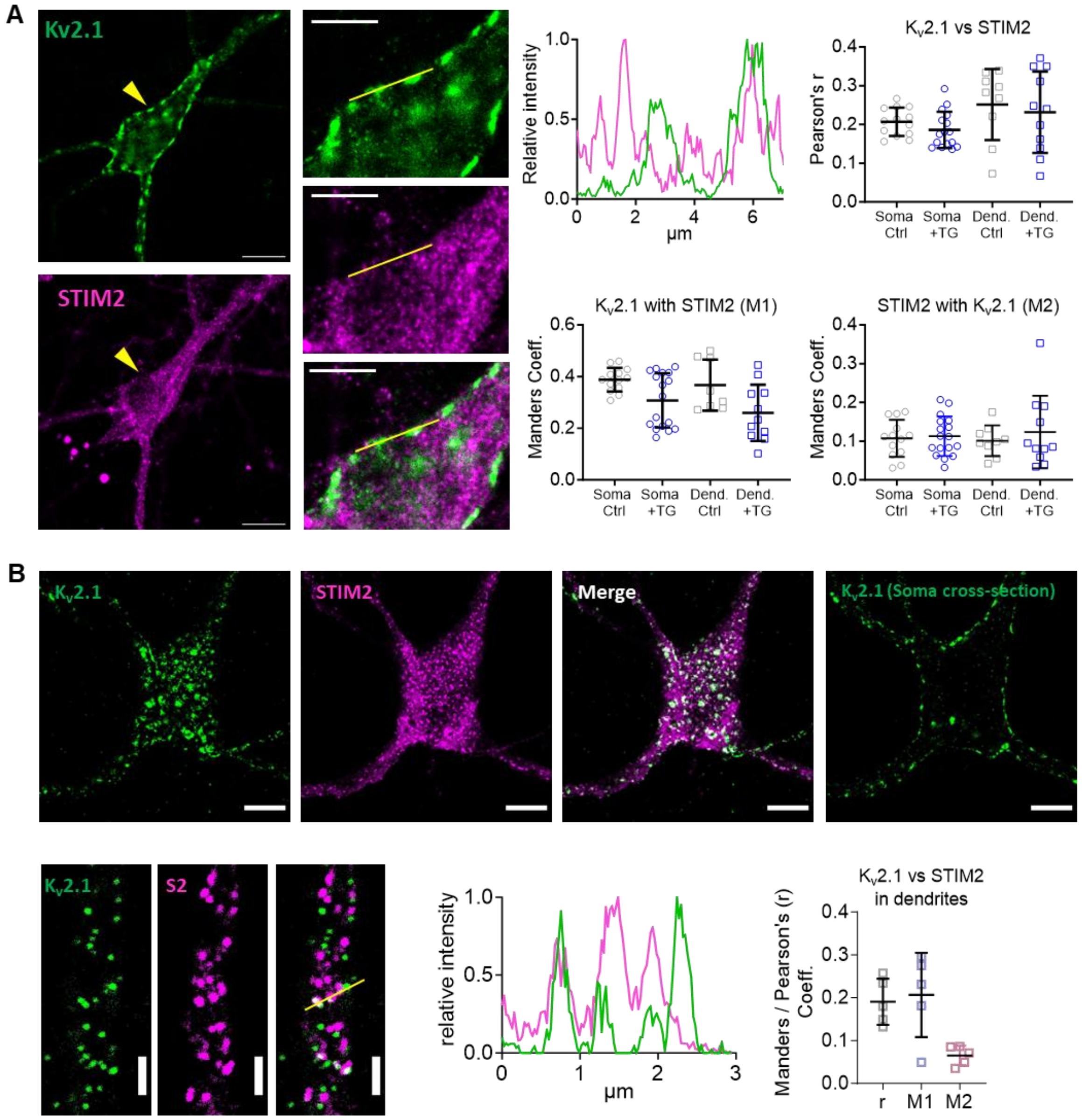

**Sup. Fig.6: A)** Colocalization analysis of Kv2.1 and STIM2 in hippocampal neurons. Immunolabeling of Kv2.1 (green) and STIM2 (magenta) in hippocampal neurons scale (bar: 20 µm) at DIV 14 under control conditions with an enlarged view (scale bar: 5 µm) of the highlighted region for the line plot in on plasma membrane. **B)** Immunolabeling of hippocampal neurons at DIV 14, showing Kv2.1 (green) and overexpressed Halo-STIM2 (magenta) under control conditions (scale bar: 10 µm), followed by soma cross-sections illustrating Kv2.1 labelling, highlighting its presence as large clusters on the plasma membrane (scale bar: 10 µm). Segment of the proximal dendrite showing scattered Kv2.1 and STIM2 clusters, with highlighting the visually overlapping spots in dendritic regions for the line plot, Scale bar: 2 µm.

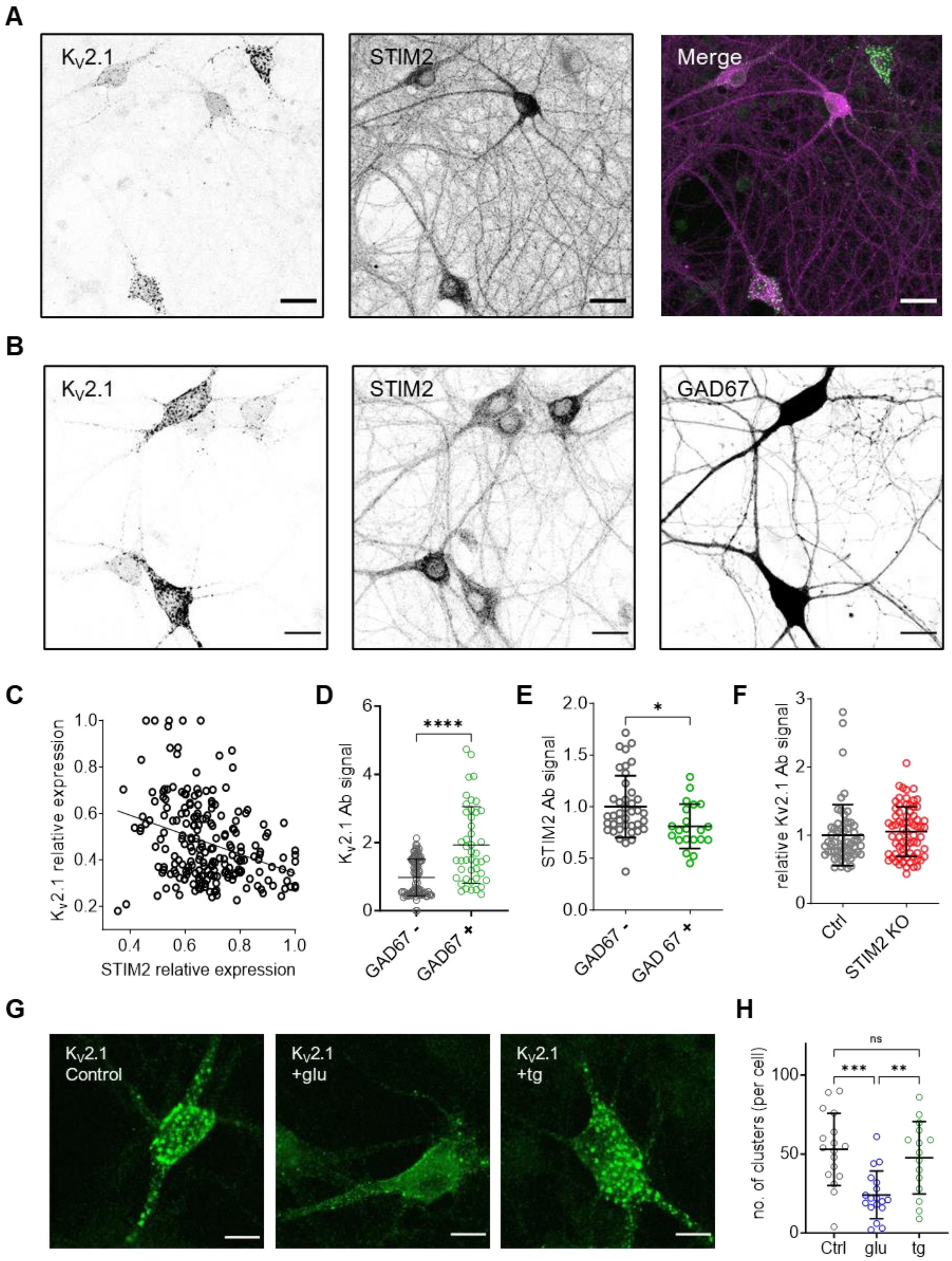

**Sup. Fig.7:** Interplay of Kv2.1 and STIM2 expression **A)** Representative immunolabeling of Kv2.1 (green) and STIM2 (magenta) in hippocampal neurons, scale bar: 20 µm. **B)** Immunolabeling of Kv2.1 and endogenous STIM2 in GAD67-GFP positive neurons from the hippocampus, stained on DIV 14-16. Scale bar: 20 µm. **C)** Scatter plot illustrating the relative expression levels of Kv2.1 and STIM2 represented by fluorescence intensity signals. **D)** Kv2.1 expression quantified from antibody signals in GAD67-positive and GAD67-negative neurons. **E)** STIM2 expression quantified in GAD67-positive and GAD67-negative neurons. Error bars denote mean ± standard deviation. **F)** Assessment of Kv2.1 expression through antibody labelling in control and Cre-mediated STIM2 knockout (KO) in hippocampal neurons derived from floxed STIM2 mice. **G)** Representative images of Kv2.1 clusters in control conditions (left), upon TG-induced ER store depletion (middle) and after NMDA treatment (right), scale bar: 10 µm **H)** Quantification of the number of clusters in resting, TG- and glutamate-stimulated cells. Quantified from Icy bioimage analysis software, version 2.4.3.0. with spot detector scale 4 (14 pixels and a 40 threshold). Refer to supplementary table for p-values.

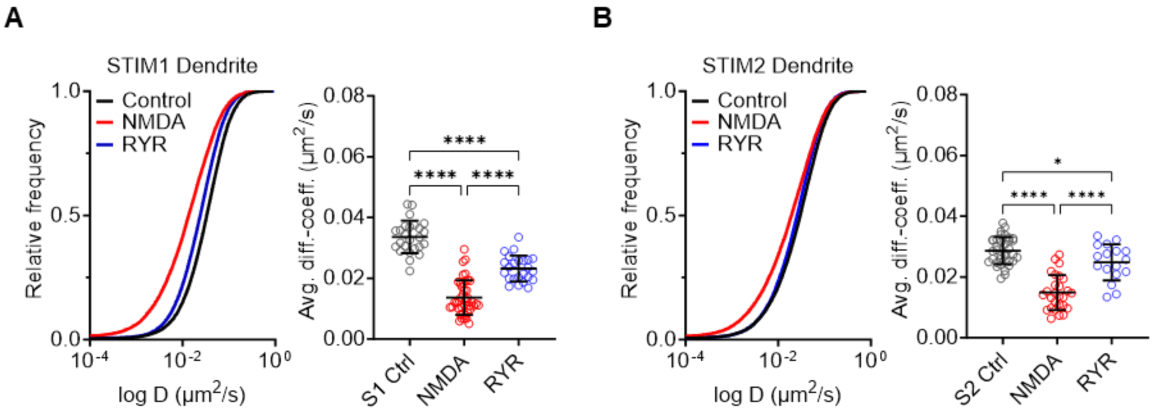

**Sup. Fig.8: A, B**) Diffusion of STIM1 (**A**) and STIM2 (**B**) in dendrites upon ryanodine (RYR) treatment. Frequency distribution and statistical comparison with NMDA treatment (1-minute NMDA exposure in $Mg^{2+}$-free Ringer's solution) and control conditions are presented as described above (with N=3 for all conditions). Acquisitions were performed in Ringer's solution containing $Mg^{2+}$ for all conditions. For the RYR experiments, ryanodine was included in the imaging medium. Statistical significance is denoted as ****$P < 0.0001$, *$P < 0.1$, determined using one-way ANOVA followed by Tukey's multiple comparisons test. Refer to supplementary table for p-values, N, n, and trajectory counts.

### A: STIM1 qRT-PCR info

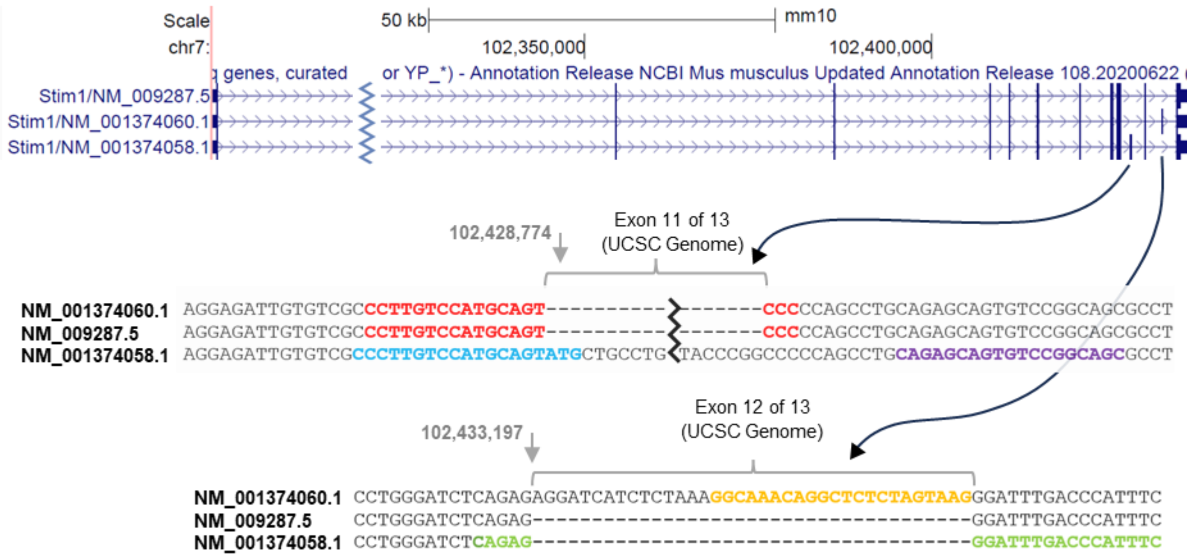

| Forward Primer | Reverse Primer | Product Amplification | Amplicon Size |
|---|---|---|---|
| CCCTTGTCCATGCAGTATG (STIM1_F_2) | GCTGCCGGACACTGCTCTG (STIM1_R_2) | NM_001374058.1 | 138 bp |
| CCTTGTCCATGCAGTCCC (STIM1_F_1_3) | GGAATGGGTCAAATCCCTCTG (STIM1_R_1_2) | NM_009287.5 | 98 bp |
| CCTTGTCCATGCAGTCCC (STIM1_F_1_3) | CTTACTAGAGAGCCTGTTTGCC (STIM1_R_3) | NM_001374060.1 | 119 bp |
| CCATGCCAAGGCTAGCAG (STIM_1const_F) | GCCGAGTCAAGAGAGGAGG (STIM_1const_R) | All 3 STIM1 Variants | 135 bp |

### B: STIM2 qRT-PCR info

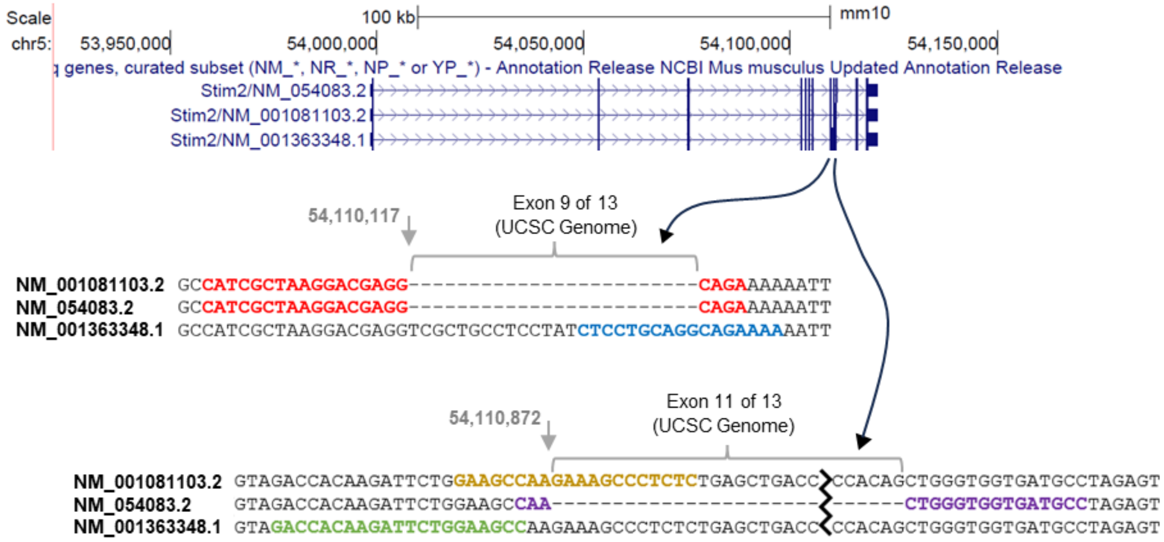

| Forward Primer | Reverse Primer | Product Amplification | Amplicon Size |
|---|---|---|---|
| CATCGCTAAGGACGAGGCAGA (STIM2-F-2-3) | GAGAGGGCTTTCTTGGCTTC (STIM2-R-1-2) | NM_001081103.2 | 129 bp |
| CATCGCTAAGGACGAGGCAGA (STIM2-F-2-3) | GGCATCACCACCCAGTTG (STIM2-R-3 new) | NM_054083.2 | 132 bp |
| CTCCTGCAGGCAGAAAAA (STIM2-F-1_new) | GGCTTCCAGAATCTTGTGGTC (STIM2-R-1) | NM_001363348.1 | 108 bp |
| CACTCCCCAGGATAGCAGTT (STIM_2const_F) | TCCTTCATCCAGTTATGAGGTG (STIM_2const_R) | All 3 STIM2 variants | 151 |

**Sup. Fig.9:**
Overview of mouse Stim1 (**A**) and Stim2 (**B**) Splice Variants and locations of designed qRTPCR primers and amplicon size on UCSE genome browser.

### Supplementary tables

Legends:
t- Total number to technical replicates. (3 technical replicate from each animal).
N- No. of animals used (for qRT-PCR) / No. of Cultures (for SPT and Staining).
n - No. of Acquisitions.
One-way ANOVA- Tukey's multiple comparisons follow-up test.
Ctrl = Control, S1= STIM1, S2= STIM2, S1Δk refers to STIM1Δk (if used).

| Fig.1 | Condition | Trajectories | n | N | Mean | SD |
|---|---|---|---|---|---|---|
| **C** | STIM1 Dendrite | 30517 | 38 | 5 | N/A | N/A |
| **C** | STIM1 Axon | 15362 | 42 | 5 | N/A | N/A |
| **D** | STIM1 Dendrite | 30517 | 38 | 5 | 0.029 | 0.0064 |
| **D** | STIM1 Axon | 15362 | 42 | 5 | 0.044 | 0.0104 |
| **D** | STIM1 Dendrite +CPA | 22709 | 32 | 4 | 0.019 | 0.0058 |
| **D** | STIM1 Axon +CPA | 4198 | 19 | 4 | 0.038 | 0.0073 |
| **F** | STIM2 Dendrite | 15133 | 24 | 3 | N/A | N/A |
| **F** | STIM2 Axon | 5635 | 26 | 3 | N/A | N/A |
| **G** | STIM2 dendrite | 15133 | 24 | 3 | 0.028 | 0.0077 |
| **G** | STIM2 Axon | 5635 | 26 | 3 | 0.058 | 0.0105 |
| **G** | STIM2 Dendrite +CPA | 13733 | 20 | 3 | 0.025 | 0.0077 |
| **G** | STIM2 Axon +CPA | 3435 | 17 | 3 | 0.052 | 0.0123 |
| **H** | STIM1 Axon | 15362 | 42 | 5 | 0.044 | 0.0104 |
| **H** | STIM2 Axon | 5635 | 26 | 3 | 0.058 | 0.0105 |
| **H** | STIM1 Dendrite | 23153 | 30 | 5 | 0.030 | 0.0045 |
| **H** | STIM2 Dendrite | 18321 | 26 | 5 | 0.026 | 0.0068 |
| **I** | Halo-TMC Dendrite | 2315 | 24 | 3 | 0.063 | 0.0062 |
| **I** | Halo-TMC Axon | 3077 | 19 | 3 | 0.066 | 0.0107 |
| **J** | STIM1 KI | 1973 | 18 | 2 | 0.024 | 0.0062 |
| **J** | STIM2 KI | 1570 | 22 | 2 | 0.019 | 0.0075 |
| **K** | STIM1 Ctrl | 13556 | 30 | 2 | 0.031 | 0.0082 |
| **K** | STIM1 KO-S1 | 15803 | 18 | 2 | 0.034 | 0.0102 |
| **K** | STIM1 KO-S2 | 44155 | 49 | 4 | 0.026 | 0.0050 |
| **K** | STIM2 Ctrl | 40603 | 39 | 4 | 0.026 | 0.0068 |
| **K** | STIM2 KO-S2 | 36027 | 47 | 4 | 0.023 | 0.0070 |

| Fig.1 | Comparison | Significance | p-value | Statistical method |
|---|---|---|---|---|
| **D** | Dendrite vs. Axon | **** | <0.0001 | ANOVA Tukey's |
| **D** | Dendrite vs. Dendrite +CPA | **** | <0.0001 | ANOVA Tukey's |
| **D** | Dendrite vs. Axon +CPA | **** | <0.0001 | ANOVA Tukey's |
| **D** | Axon vs. Dendrite +CPA | **** | <0.0001 | ANOVA Tukey's |
| **D** | Axon vs. Axon +CPA | ns | 0.1093 | ANOVA Tukey's |
| **D** | Dendrite + CPA vs. Axon +CPA | **** | <0.0001 | ANOVA Tukey's |
| **G** | Dendrite vs. Axon | **** | <0.0001 | ANOVA Tukey's |
| **G** | Dendrite vs. Dendrite +CPA | ns | 0.7393 | ANOVA Tukey's |
| **G** | Dendrite vs. Axon +CPA | **** | <0.0001 | ANOVA Tukey's |
| **G** | Axon vs. Dendrite +CPA | **** | <0.0001 | ANOVA Tukey's |
| **G** | Axon vs. Axon +CPA | ns | 0.2750 | ANOVA Tukey's |
| **G** | Dendrite + CPA vs. Axon +CPA | **** | <0.0001 | ANOVA Tukey's |
| **H** | STIM1 vs. STIM2 (Axons) | **** | <0.0001 | Unpaired t test |
| **H** | STIM1 vs. STIM2 (Dendrites) | * | 0.0149 | Unpaired t test |
| **I** | Halo-TMC Axon vs. Dendrite | ns | 0.326 | Unpaired t test |
| **J** | STIM1 KI vs. STIM2 KI | * | 0.035 | Unpaired t test |
| **K** | STIM1 Ctrl vs. KO-S1 | ns | >0.9999 | ANOVA Tukey's |
| **K** | STIM1 Ctrl vs. KO-S2 | * | 0.0312 | ANOVA Tukey's |
| **K** | STIM1 KO-S1 vs. KO-S2 | ** | 0.0076 | ANOVA Tukey's |
| **K** | STIM2 Ctrl vs. KO-S2 | * | 0.0293 | Unpaired t test |

| Fig.2 | Condition | Trajectories | n | N | Mean | SD |
|---|---|---|---|---|---|---|
| **B-D** | STIM1 Ctrl | N/A | 07 | 2 | N/A | N/A |
| **B-D** | STIM2 TG | N/A | 03 | 2 | N/A | N/A |
| **B-D** | STIM2 Ctrl | N/A | 04 | 2 | N/A | N/A |
| **B-D** | STIM2 TG | N/A | 05 | 2 | N/A | N/A |
| **F** | STIM1 Syn-ROI | 2235 | 27 | 2 | 0.027 | 0.0066 |

| | | | | | | |
|---|---|---|---|---|---|---|
| F | STIM1 Extra-synaptic | 6859 | 28 | 2 | 0.026 | 0.0077 |
| F | STIM1 Syn-ROI +CPA | 1299 | 12 | 2 | 0.024 | 0.0042 |
| F | STIM1 Extra-synaptic +CPA | 4337 | 11 | 2 | 0.023 | 0.0045 |
| H | STIM2 Syn-ROI | 1016 | 21 | 2 | 0.027 | 0.0077 |
| H | STIM2 Extra-synaptic | 3863 | 22 | 2 | 0.032 | 0.0096 |
| H | STIM2 Syn-ROI +CPA | 1542 | 18 | 2 | 0.027 | 0.0044 |
| H | STIM2 Extra-synaptic +CPA | 5745 | 18 | 2 | 0.033 | 0.0042 |
| J | Halo-TMC Syn-ROI | 273 | 09 | 1 | 0.050 | 0.0114 |
| J | Halo-TMC Extra-synaptic | 533 | 09 | 1 | 0.053 | 0.0095 |

| Fig.2 | Comparison | Significance | p-value | Statistical method |
|---|---|---|---|---|
| B | STIM1 Ctrl vs. TG | ns | 0.29 | Unpaired t test |
| B | STIM2 Ctrl vs. TG | ns | 0.13 | Unpaired t test |
| C | STIM1 Ctrl vs. TG | ns | 0.27 | Unpaired t test |
| C | STIM2 Ctrl vs. TG | ns | 0.31 | Unpaired t test |
| D | STIM1 Ctrl vs. TG | *** | 0.0007 | Unpaired t test |
| D | STIM2 Ctrl vs. TG | ns | 0.56 | Unpaired t test |
| F | STIM1 syn vs. ex-syn | ns | 0.9894 | ANOVA Tukey's |
| F | STIM1 syn vs. syn +CPA | ns | 0.7005 | ANOVA Tukey's |
| F | STIM1 syn vs. ex-syn +CPA | ns | 0.2682 | ANOVA Tukey's |
| F | STIM1 ex-syn vs. syn +CPA | ns | 0.8322 | ANOVA Tukey's |
| F | STIM1 ex-syn vs. ex-syn +CPA | ns | 0.3837 | ANOVA Tukey's |
| F | STIM1 syn +CPA vs. ex-syn +CPA | ns | 0.9093 | ANOVA Tukey's |
| H | STIM2 syn vs. ex-syn | ns | 0.0934 | ANOVA Tukey's |
| H | STIM2 syn vs. syn +CPA | ns | 0.9995 | ANOVA Tukey's |
| H | STIM2 syn vs. ex-syn +CPA | * | 0.0416 | ANOVA Tukey's |
| H | STIM2 ex-syn vs. syn +CPA | ns | 0.0899 | ANOVA Tukey's |
| H | STIM2 ex-syn vs. ex-syn +CPA | ns | 0.9670 | ANOVA Tukey's |
| H | STIM2 syn +CPA vs. ex-syn +CPA | * | 0.0407 | ANOVA Tukey's |
| J | STIM-TMC syn-ROI vs. ex-syn | ns | 0.5217 | Unpaired t test |

| Fig.3 | Condition | n | N |
|---|---|---|---|
| E-G | STIM1 Ctrl | 07 | 2 |
| E-G | STIM2 TG | 09 | 2 |
| E-G | STIM2 Ctrl | 06 | 2 |
| E-G | STIM2 TG | 09 | 2 |

| Fig.3 | Comparison | Significance | p-value | Statistical method |
|---|---|---|---|---|
| E | STIM1 Ctrl vs. TG | ns | 0.14 | Unpaired t test |
| E | STIM2 Ctrl vs. TG | ns | 0.99 | Unpaired t test |
| F | STIM1 Ctrl vs. TG | ns | 0.10 | Unpaired t test |
| F | STIM2 Ctrl vs. TG | ns | 0.82 | Unpaired t test |
| G | STIM1 Ctrl vs. TG | ns | 0.62 | Unpaired t test |
| G | STIM2 Ctrl vs. TG | * | 0.02 | Unpaired t test |

| Fig.4 | Condition | Trajectories | n | N | Mean | SD |
|---|---|---|---|---|---|---|
| B | STIM1 Control | 16280 | 19 | 3 | N/A | N/A |
| B | STIM1 glu+APV | 16543 | 17 | 3 | N/A | N/A |
| B | STIM1 +glu | 15409 | 19 | 3 | N/A | N/A |
| C | STIM1 Control | 16280 | 19 | 3 | 0.032 | 0.0064 |
| C | STIM1 glu+APV | 16543 | 17 | 3 | 0.013 | 0.0074 |
| C | STIM1 +glu | 15409 | 19 | 3 | 0.028 | 0.0078 |
| E | STIM2 Control | 18696 | 18 | 3 | N/A | N/A |
| E | STIM2 glu+APV | 11725 | 19 | 3 | N/A | N/A |
| E | STIM2 +glu | 8045 | 15 | 3 | N/A | N/A |
| F | STIM2 Control | 18696 | 18 | 3 | 0.027 | 0.0051 |
| F | STIM2 glu+APV | 11725 | 19 | 3 | 0.015 | 0.0063 |
| F | STIM2 +glu | 8045 | 15 | 3 | 0.024 | 0.0077 |
| H | STIM1Δk Dendrite Control | 12013 | 11 | 2 | 0.035 | 0.0025 |
| H | STIM1Δk Dendrite +glu | 13160 | 15 | 2 | 0.024 | 0.0105 |
| I | STIM1 +MK-801 | 27777 | 27 | 2 | 0.033 | 0.0061 |
| I | STIM2 Control | 27595 | 35 | 2 | 0.028 | 0.0045 |
| I | STIM2 +MK-801 | 23765 | 36 | 2 | 0.034 | 0.0081 |

| | | | | | | |
|---|---|---|---|---|---|---|
| **M** | CPA | N/A | 6 | 2 | 0.49 | 0.24 |
| **M** | NMDA | N/A | 6 | 2 | 0.46 | 0.12 |

| Fig.4 | Comparison | Significance | p-value | Statistical method |
|---|---|---|---|---|
| **C** | STIM1 Control vs. glutamate | **** | <0.0001 | ANOVA Tukey's |
| **C** | STIM1 Control vs. glu + APV | ns | 0.2356 | ANOVA Tukey's |
| **C** | STIM1 glutamate vs. glu + APV | **** | <0.0001 | ANOVA Tukey's |
| **F** | STIM2 Control vs. glutamate | **** | <0.0001 | ANOVA Tukey's |
| **F** | STIM2 Control vs. glu + APV | ns | 0.3579 | ANOVA Tukey's |
| **F** | STIM2 glutamate vs. glu + APV | *** | 0.0003 | ANOVA Tukey's |
| **H** | S1 Ctrl vs. S1 +glu | **** | <0.0001 | ANOVA Tukey's |
| **H** | S1 Ctrl vs. S1 +glu +APV | ns | 0.5809 | ANOVA Tukey's |
| **H** | S1 Ctrl vs. S1Δk Ctrl | ns | 0.826 | ANOVA Tukey's |
| **H** | S1 Ctrl vs. S1Δk +glu | * | 0.0249 | ANOVA Tukey's |
| **H** | S1 +glu vs. S1 +glu +APV | **** | <0.0001 | ANOVA Tukey's |
| **H** | S1 +glu vs. S1Δk Ctrl | **** | <0.0001 | ANOVA Tukey's |
| **H** | S1 +glu vs. S1Δk +glu | ** | 0.0023 | ANOVA Tukey's |
| **H** | S1 +glu +APV vs. S1Δk Ctrl | ns | 0.1484 | ANOVA Tukey's |
| **H** | S1 +glu +APV vs. S1Δk +glu | ns | 0.4575 | ANOVA Tukey's |
| **H** | S1Δk Ctrl vs. S1Δk +glu | ** | 0.0039 | ANOVA Tukey's |
| **I** | S1 Ctrl vs. S1 +MK801 | ns | 0.9995 | ANOVA Tukey's |
| **I** | S1 Ctrl vs. S2 Ctrl | * | 0.0166 | ANOVA Tukey's |
| **I** | S1 Ctrl vs. S2 +MK801 | ns | 0.8393 | ANOVA Tukey's |
| **I** | S1 +MK801 vs. S2 Ctrl | * | 0.0192 | ANOVA Tukey's |
| **I** | S1 +MK801 vs. S2 +MK801 | ns | 0.7672 | ANOVA Tukey's |
| **I** | S2 Ctrl vs. S +MK801 | *** | 0.0003 | ANOVA Tukey's |
| **K** | S1Ctrl vs. S1TG | **** | <0.0001 | Kruskal-Wallis, Dunn's |
| **K** | S1Ctrl vs. S1NMDA | **** | <0.0001 | Kruskal-Wallis, Dunn's |
| **K** | S1TG vs. S1NMDA | **** | <0.0001 | Kruskal-Wallis, Dunn's |
| **K** | S2Ctrl vs. S2TG | **** | <0.0001 | Kruskal-Wallis, Dunn's |
| **K** | S2Ctrl vs. S2NMDA | **** | <0.0001 | Kruskal-Wallis, Dunn's |
| **K** | S2TG vs. S2NMDA | * | 0.0181 | Kruskal-Wallis, Dunn's |
| **M** | Amp. CPA vs. NMDA | ns | 0.7886 | Unpaired t test |
| **M** | Tau (s) CPA vs. NMDA | *** | 0.002 | Unpaired t test |

| Fig.5 | Condition | Trajectories | n | N | Mean | SD |
|---|---|---|---|---|---|---|
| **B** | STIM1 Dendrite Control | 23153 | 30 | 5 | 0.030 | 0.0045 |
| **B** | STIM1 Dendrite +NMDA | 12995 | 25 | 5 | 0.015 | 0.0061 |
| **B** | STIM1 Dendrite +NMDA+nim | 25471 | 46 | 5 | 0.017 | 0.0052 |
| **C** | ER calcium- NMDA | N/A | 7 | 2 | N/A | N/A |
| **C** | ER calcium- NMDA + nim | N/A | 6 | 2 | N/A | N/A |
| **D** | STIM1 Dendrite Control | 1147 | 08 | 1 | 0.023 | 0.0027 |
| **D** | STIM1 NMDA (1-5min) | 1544 | 05 | 1 | 0.013 | 0.0050 |
| **D** | STIM1 NMDA (5-10min) | 2221 | 05 | 1 | 0.025 | 0.0048 |
| **E** | STIM1 Dendrite Control | 1147 | 08 | 1 | 0.023 | 0.0027 |
| **E** | STIM1 NMDA + nim (1-5min) | 1335 | 05 | 1 | 0.015 | 0.0022 |
| **E** | STIM1 NMDA + nim (5-10min) | 1505 | 04 | 1 | 0.021 | 0.0022 |
| **G** | STIM2 Dendrite Control | 24814 | 32 | 4 | 0.026 | 0.0076 |
| **G** | STIM2 Dendrite +NMDA | 18169 | 21 | 4 | 0.015 | 0.0054 |
| **G** | STIM2 Dendrite +NMDA+nim | 15623 | 18 | 4 | 0.014 | 0.0056 |

| Fig.5 | Comparison | Significance | p-value | Statistical method |
|---|---|---|---|---|
| **B** | STIM1 Control vs. NMDA | **** | <0.0001 | ANOVA Tukey's |
| **B** | STIM1 Control vs. nim+NMDA | **** | <0.0001 | ANOVA Tukey's |
| **B** | STIM1 NMDA vs. nim+NMDA | ns | 0.2228 | ANOVA Tukey's |
| **C** | NMDA vs. nim+NMDA | ns | 0.6800 | Unpaired t-test |
| **G** | STIM2 Control vs. NMDA | **** | <0.0001 | ANOVA Tukey's |
| **G** | STIM2 Control vs. nim+NMDA | **** | <0.0001 | ANOVA Tukey's |
| **G** | STIM2 NMDA vs. nim+NMDA | ns | 0.8970 | ANOVA Tukey's |
| **I** | STIM1 Knockout vs. Δ CRE (basal $Ca^{2+}$) | ns | >0.999 | Unpaired t-test |
| **I** | STIM1 Knockout vs. Δ CRE ($K^+$ induced influx) | ns | 0.6737 | Unpaired t-test |
| **J** | STIM2 Knockout vs. Δ CRE (basal $Ca^{2+}$) | ns | >0.999 | Unpaired t-test |
| **J** | STIM2 Knockout vs. Δ CRE ($K^+$ induced influx) | ns | 0.7665 | Unpaired t-test |
| **K** | TG vs. DMSO peak current | ns | 0.9254 | Unpaired t-test |
| **L** | STIM1 over expressed vs. wildtype peak current | ns | 0.5544 | Unpaired t-test |

| Fig.6 | Condition | Trajectories | n | N | Mean | SD |
|---|---|---|---|---|---|---|
| **B-D** | STIM1 Ctrl | N/A | 18 | 3 | N/A | N/A |
| **B-D** | STIM2 TG | N/A | 15 | 3 | N/A | N/A |
| **B-D** | STIM2 Ctrl | N/A | 18 | 3 | N/A | N/A |
| **B-D** | STIM2 TG | N/A | 20 | 3 | N/A | N/A |
| **G** | $K_V$2.1 | detected clusters | = 270 | 2 | 2.74 | 1.29 |
| **G** | MAPPER | detected clusters | = 3330 | 2 | 0.57 | 0.21 |
| **I** | STIM1 in MAPPER Control | 836 | 07 | 2 | 0.018 | 0.0071 |
| **I** | STIM1 in MAPPER +NMDA | 368 | 08 | 2 | 0.007 | 0.0040 |
| **K** | STIM2 in MAPPER Control | 205 | 10 | 2 | 0.0062 | 0.0031 |
| **K** | STIM2 in MAPPER +NMDA | 370 | 08 | 2 | 0.0055 | 0.0010 |

| Fig.6 | Comparison | Significance | p-value | Statistical method |
|---|---|---|---|---|
| **B** | STIM1 Ctrl vs. TG | ns | 0.46 | Unpaired t test |
| **B** | STIM2 Ctrl vs. TG | * | 0.02 | Unpaired t test |
| **C** | STIM1 Ctrl vs. TG | ns | 0.99 | Unpaired t test |
| **C** | STIM2 Ctrl vs. TG | ns | 0.16 | Unpaired t test |
| **D** | STIM1 Ctrl vs. TG | ns | 0.52 | Unpaired t test |
| **D** | STIM2 Ctrl vs. TG | ns | 0.87 | Unpaired t test |
| **G** | $K_V$2.1 vs. MAPPER | **** | <0.0001 | Unpaired t test |
| **I** | STIM1 Control vs. NMDA | ** | 0.0023 | Unpaired t test |
| **K** | STIM2 Control vs. NMDA | ns | 0.5559 | Unpaired t test |

**Table S1:** Statistical data corresponding to qRT-PCR

| Sup. Fig. 1C: All Stim1 variants normalized to constitutive Stim1 expression. | | | | | | | | | |
|---|---|---|---|---|---|---|---|---|---|
| | Stim1s | | | Stim1l | | | Stim1b | | |
| **Region** | Cortex | Hippo. | Cereb. | Cortex | Hippo. | Cereb. | Cortex | Hippo. | Cereb. |
| **Mean** | 0.214 | 0.289 | 0.133 | 0.074 | 0.065 | 0.080 | 0.062 | 0.040 | 0.078 |
| **SD** | 0.042 | 0.080 | 0.034 | 0.056 | 0.039 | 0.052 | 0.028 | 0.018 | 0.020 |
| **SEM** | 0.017 | 0.032 | 0.013 | 0.023 | 0.016 | 0.021 | 0.011 | 0.007 | 0.008 |
| **t/N** | 6/2 | 6/2 | 6/2 | 6/2 | 6/2 | 6/2 | 6/2 | 6/2 | 6/2 |

| Sup. Fig. 1D: Stim1s normalized with STIM2.2. | | | | | | |
|---|---|---|---|---|---|---|
| | Stim1s | | | Stim2.2 | | |
| **Region** | Cortex | Hippo. | Cereb. | Cortex | Hippo. | Cereb. |
| **Mean** | 0.145 | 0.071 | 0.043 | 1.002 | 1.008 | 1.003 |
| **SD** | 0.032 | 0.031 | 0.024 | 0.065 | 0.144 | 0.078 |
| **SEM** | 0.013 | 0.012 | 0.010 | 0.026 | 0.059 | 0.032 |
| **t/N** | 6/2 | 6/2 | 6/2 | 6/2 | 6/2 | 6/2 |

| Sup Fig. 1E: All Stim2 variants normalized to constitutive Stim2 expression. | | | | | | | | | |
|---|---|---|---|---|---|---|---|---|---|
| | Stim2.1 | | | Stim2.2 | | | Stim2.2 | | |
| **Region** | Cortex | Hippo. | Cereb. | Cortex | Hippo. | Cereb. | Cortex | Hippo. | Cereb. |
| **Mean** | 0.018 | 0.016 | 0.026 | 1.479 | 1.526 | 1.223 | 0.013 | 0.009 | 0.026 |
| **SD** | 0.018 | 0.018 | 0.004 | 0.165 | 0.246 | 0.157 | 0.008 | 0.005 | 0.018 |
| **SEM** | 0.007 | 0.007 | 0.002 | 0.067 | 0.100 | 0.064 | 0.003 | 0.002 | 0.009 |
| **t/N** | 6/2 | 6/2 | 3/2 | 6/2 | 6/2 | 6/2 | 6/2 | 4/2 | 4/2 |

| Sup. Fig.1F: Stim1 Cons normalized with Stim2 Cons. | | | | | | |
|---|---|---|---|---|---|---|
| | Stim1 Constitutive | | | Stim2 Constitutive | | |
| **Region** | Cortex | Hippo. | Cereb. | Cortex | Hippo. | Cereb. |
| **Mean** | 1.033 | 0.391 | 0.670 | 1.007 | 1.002 | 1.002 |
| **SD** | 0.306 | 0.185 | 0.133 | 0.130 | 0.069 | 0.075 |
| **SEM** | 0.124 | 0.075 | 0.054 | 0.053 | 0.028 | 0.030 |
| **t/N** | 6/2 | 6/2 | 6/2 | 6/2 | 6/2 | 6/2 |

**Table S2:**

| Sup. Fig. | Comparison | Significance | p-value | Statistical method |
|---|---|---|---|---|
| **3A** | STIM1 vs. Halo-TMC | *** | 0.0002 | ANOVA Tukey's |
| **3A** | STIM1 vs. STIM2 | ns | 0.9036 | ANOVA Tukey's |

| | | | | |
|---|---|---|---|---|
| **3A** | Halo-TMC vs. STIM2 | *** | 0.0001 | ANOVA Tukey's |
| **3B** | STIM1 vs. Halo-TMC | **** | <0.0001 | ANOVA Tukey's |
| **3B** | STIM1 vs. STIM2 | ns | 0.5958 | ANOVA Tukey's |
| **3B** | Halo-TMC vs. STIM2 | ** | 0.0013 | ANOVA Tukey's |
| **5B** | STIM1 Cre (-) vs. Cre (+) | **** | <0.0001 | Unpaired t-test |
| **5D** | STIM2 Cre (-) vs. Cre (+) | **** | <0.0001 | Unpaired t-test |
| **7C** | Pearson’s r = -0.3239, R2 = 0.1049 | N/A | N/A | N/A |
| **7D** | GAD- vs. GAD+ | **** | <0.0001 | Unpaired t-test |
| **7E** | GAD- vs. GAD+ | * | 0.0123 | Unpaired t-test |
| **7F** | Control vs STIM2 KD | ns | 0.4629 | Unpaired t-test |
| **7H** | Ctrl vs. glu | *** | 0.0004 | ANOVA Tukey's |
| **7H** | Ctrl vs. tg | ns | 0.7423 | ANOVA Tukey's |
| **7H** | glu vs. tg | ** | 0.0054 | ANOVA Tukey's |
| **8A** | S1 Ctrl vs. NMDA | **** | <0.0001 | ANOVA Tukey's |
| **8A** | S1 Ctrl vs. RYR | **** | <0.0001 | ANOVA Tukey's |
| **8A** | NMDA vs. RYR | **** | <0.0001 | ANOVA Tukey's |
| **8B** | S2 Ctrl vs. NMDA | **** | <0.0001 | ANOVA Tukey's |
| **8B** | S2 Ctrl vs. RYR | * | 0.0422 | ANOVA Tukey's |
| **8B** | NMDA vs. RYR | **** | <0.0001 | ANOVA Tukey's |

| **Fig.Sup.4** | Condition | Trajectories | n | N | Mean | SD |
|---|---|---|---|---|---|---|
| **A** | STIM1 dendrite Control | 40417 | 24 | 3 | 0.026 | 0.0057 |
| **A** | STIM1 dendrite BIC | 24567 | 23 | 3 | 0.019 | 0.0051 |
| **A** | STIM1 dendrite TTX | 29544 | 25 | 3 | 0.033 | 0.0053 |
| **A** | STIM1 Dendrite Control | 41818 | 25 | 2 | 0.0287 | 0.0048 |
| **A** | STIM1 Dendrite DHPG | 17264 | 15 | 2 | 0.0268 | 0.0049 |
| **B** | STIM2 dendrite Control | 13217 | 18 | 3 | 0.033 | 0.0117 |
| **B** | STIM2 dendrite BIC | 20966 | 23 | 3 | 0.031 | 0.0092 |
| **B** | STIM2 dendrite TTX | 14140 | 24 | 3 | 0.028 | 0.0089 |
| **B** | STIM2 Dendrite Control | 14205 | 17 | 2 | 0.031 | 0.0093 |
| **B** | STIM2 Dendrite DHPG | 8926 | 11 | 2 | 0.028 | 0.0077 |
| **C** | STIM1 dendrite Control | 2334 | 18 | 2 | 0.026 | 0.0075 |
| **C** | STIM1 dendrite BayK | 3176 | 19 | 2 | 0.027 | 0.0069 |
| **D** | STIM1 Axon Control | 7267 | 12 | 2 | 0.039 | 0.0054 |
| **D** | STIM1 Axon glutamate | 10012 | 12 | 2 | 0.035 | 0.0088 |
| **E** | STIM1 Axon Control | 11447 | 25 | 2 | 0.046 | 0.0085 |
| **E** | STIM1 Axon DHPG | 6138 | 15 | 2 | 0.047 | 0.0098 |
| **F** | STIM1 axon Control | 6494 | 20 | 3 | 0.046 | 0.0114 |
| **F** | STIM1 axon BIC | 5212 | 21 | 3 | 0.040 | 0.0100 |
| **F** | STIM1 axon TTX | 7034 | 22 | 3 | 0.045 | 0.0106 |
| **G** | STIM2 Axon Control | 3436 | 08 | 1 | 0.050 | 0.0086 |
| **G** | STIM2 Axon glutamate | 3117 | 07 | 1 | 0.045 | 0.0093 |
| **H** | STIM2 Axon Control | 2237 | 15 | 2 | 0.059 | 0.0110 |
| **H** | STIM2 Axon DHPG | 1236 | 06 | 2 | 0.059 | 0.0047 |
| **I** | STIM2 axon Control | 577 | 05 | 1 | 0.053 | 0.0076 |
| **I** | STIM2 axon BIC | 90 | 04 | 1 | 0.049 | 0.0048 |
| **I** | STIM2 axon TTX | 655 | 08 | 1 | 0.049 | 0.0127 |
| **J** | STIM1Δk Axon Control | 8314 | 07 | 1 | 0.040 | 0.0025 |
| **J** | STIM1Δk Axon glutamate | 8469 | 07 | 1 | 0.042 | 0.0067 |

| **Fig.Sup.4** | Comparison | Significance | p-value | Statistical method |
|---|---|---|---|---|
| **A** | STIM1 Dendrite Control vs. BIC | **** | <0.0001 | ANOVA Tukey's |
| **A** | STIM1 Dendrite Control vs. TTX | **** | <0.0001 | ANOVA Tukey's |
| **A** | STIM1 Dendrite TTX vs. BIC | **** | <0.0001 | ANOVA Tukey's |
| **A** | STIM1 Dendrite Control vs. DHPG | ns | 0.2304 | Unpaired t test |
| **B** | STIM2 Dendrite Control vs. BIC | ns | 0.6660 | ANOVA Tukey's |
| **B** | STIM2 Dendrite Control vs. TTX | ns | 0.2773 | ANOVA Tukey's |
| **B** | STIM2 Dendrite TTX vs. BIC | ns | 0.7519 | ANOVA Tukey's |
| **B** | STIM2 Dendrite Control vs. DHPG | ns | 0.3219 | Unpaired t test |
| **C** | STIM1 Dendrite Control vs. BayK | ns | 0.7968 | Unpaired t test |
| **D** | STIM1 Axon Control vs. glutamate | ns | 0.1854 | Unpaired t test |
| **E** | STIM1 Axon Control vs. DHPG | ns | 0.5089 | Unpaired t test |
| **F** | STIM1 Axon Control vs. BIC | ns | 0.1760 | ANOVA Tukey's |
| **F** | STIM1 Axon Control vs. TTX | ns | 0.9747 | ANOVA Tukey's |
| **F** | STIM1 Axon TTX vs. BIC | ns | 0.2401 | ANOVA Tukey's |
| **G** | STIM2 Axon Control vs. glutamate | ns | 0.2509 | Unpaired t test |

| | | | | |
|---|---|---|---|---|
| **H** | STIM2 Axon Control vs. DHPG | ns | 0.9309 | Unpaired t test |
| **I** | STIM2 Axon Control vs. BIC | ns | 0.9936 | ANOVA Tukey's |
| **I** | STIM2 Axon Control vs. TTX | ns | 0.9764 | ANOVA Tukey's |
| **I** | STIM2 Axon TTX vs. BIC | ns | 0.9998 | ANOVA Tukey's |
| **J** | STIM1Δk Axon Ctrl vs. glutamate | ns | 0.5167 | Unpaired t test |
